# Epidemic Growth Rates and Host Movement Patterns Shape Management Performance for Pathogen Spillover at The Wildlife-livestock Interface

**DOI:** 10.1101/617290

**Authors:** Kezia R. Manlove, Laura M. Sampson, Benny Borremans, E. Frances Cassirer, Ryan S. Miller, Kim M. Pepin, Thomas E. Besser, Paul C. Cross

**Affiliations:** Department of Wildland Resources and Ecology Center Utah State University, Logan, UT, 84321; Center for Infectious Disease Dynamics Pennsylvania State University, University Park, PA, 16802; Department of Ecology Evolutionary Biology University of California, Los Angeles, Los Angeles, California; Idaho Department of Fish and Game 3316 16th Street, Lewiston, ID, 83501; United States Department of Agriculture, Animal and Plant Health Inspection Service Center for Epidemiology and Animal Health, Fort Collins, CO, USA 80523; National Wildlife Research Center USDA-APHIS, Wildlife Services 4101 Laporte Ave., Fort Collins, CO 80521; Department of Veterinary Microbiology and Pathology Washington State University, Pullman, WA, 99164-7040; U. S. Geological Survey, Northern Rocky Mountain Science Center Bozeman, MT 59715

**Keywords:** pathogen spillover, wildlife-livestock interface, disease management, structured decision making, disease model, dispersal kernel

## Abstract

Managing pathogen spillover at the wildlife-livestock interface is a key step toward improving global animal health, food security, and wildlife conservation. However, predicting the effectiveness of management actions across host-pathogen systems with different life histories is an on-going challenge since data on intervention effectiveness are expensive to collect and results are system-specific. We developed a simulation model to explore how the efficacies of different management strategies vary according to host movement patterns and epidemic growth rates. The model suggested that fast-growing, fast-moving epidemics like avian influenza were best-managed with actions like biosecurity or containment, which limited and localized overall spillover risk. For fast-growing, slower-moving diseases like foot-and-mouth disease, depopulation or prophylactic vaccination were competitive management options. Many actions performed competitively when epidemics grew slowly and host movements were limited, and how management efficacy related to epidemic growth rate or host movement propensity depended on what objective was used to evaluate management performance. This framework may be a useful step in advancing how we classify and prioritise responses to novel pathogen spillover threats, and evaluate current management actions for pathogens emerging at the wildlife-livestock interface.

## 1 Introduction

Cross-species spillover of pathogens occurs when a pathogen that is released by a member of a reservoir host species goes on to establish and replicate in a different (recipient) host species [1, 2, 3]. While mitigating pathogen spillover and associated disease risk at the wildlife-livestock interface is a major goal for both human and animal health agencies [4], spillover management decisions are often context-specific and based on expert opinion [5]. In particular, there is limited scientific knowledge to guide management of spillover risk in understudied systems. Here, we propose a modeling framework designed to fill that gap by providing evidence-based guidance about optimal management over a wide range of ecological contexts.

A pathogen’s spatial extent, and the spatial connectivity in the host populations, are critical drivers of disease management efficacy for many important wildlife-livestock [6] and wildlife-human spillover systems [6, 7, 8], and management of reservoir versus recipient hosts has been compared in some contexts [9]. However, spatially explicit two-host disease models have received less attention, with most published models describing management efficacy within a single host species [11]. Moreover, these single-host models are often not intended to address spillover risk per se, but rather, to characterise dynamics leading up to or trailing after a spillover event.

Another group of models builds on the idea that spillover disease burdens depend on both the frequency and the consequences of individual spillover events. These “multi-host” models often compare within- and between-species transmission rates to identify pathogens with a high risk of generating catastrophic spillover events [10, 11, 12]. Multi-host models have proven useful for characterising spillover rates, especially when merged with phylogenetic information specifically identifying the source of a given spillover event [15]. However, they are rarely extended to account for changing spillover risk as local reservoir prevalences vary through space and time.

Here, we explore how to best manage pathogen spillover and transmission in wildlife-livestock systems, using actions that are spatially explicit and could be applied to either the reservoir or the recipient host species. We start by justifying why a management-centered framework for wildlife-livestock spillover should explicitly incorporate movement. We then go on to describe a simple disease propagation model to characterise spillover risk and onward transmission across a wide range of hosts and pathogens. We next use the model to explore spillover management efficacies, and describe the patterns that the model produced. We end by discussing the limitations of this approach, and outlining features that could be added in the future.

## 2 Characterising disease propagation in terms of epidemic growth rates and host movements

Our model is structured around three initial conjectures. First, we anticipate that the most efficient means of controlling pathogen spillover could often rest on the spatial dynamics of the hosts. Hosts with high movement propensities can produce widespread spatial synchrony in spillover risk, making the precise location of future spillover events hard to predict (**Plowright et al. *in revision***). In these cases, the best management option may be to target interspecific contacts by applying biosecurity measures or phytosanitary controls across a broad spatial extent [13]. When reservoir hosts move shorter distances, however, spatial containment in either the reservoir or the recipient host (or both) may be possible.

Second, we anticipate that the relative efficacies of various management actions will depend on how quickly an epidemic grows. If epidemic growth is rapid in the recipient host population, then depopulation or targeted vaccination at the time of the outbreak (here referred to as retroactive vaccination) may effectively limit post-spillover epidemic size. If epidemics grow slowly, actions like prophylactic vaccination of the reservoir population may lower reservoir prevalence to the point where any cross-species spillover event becomes vanishingly rare (Figure 1).

**Figure 1:**
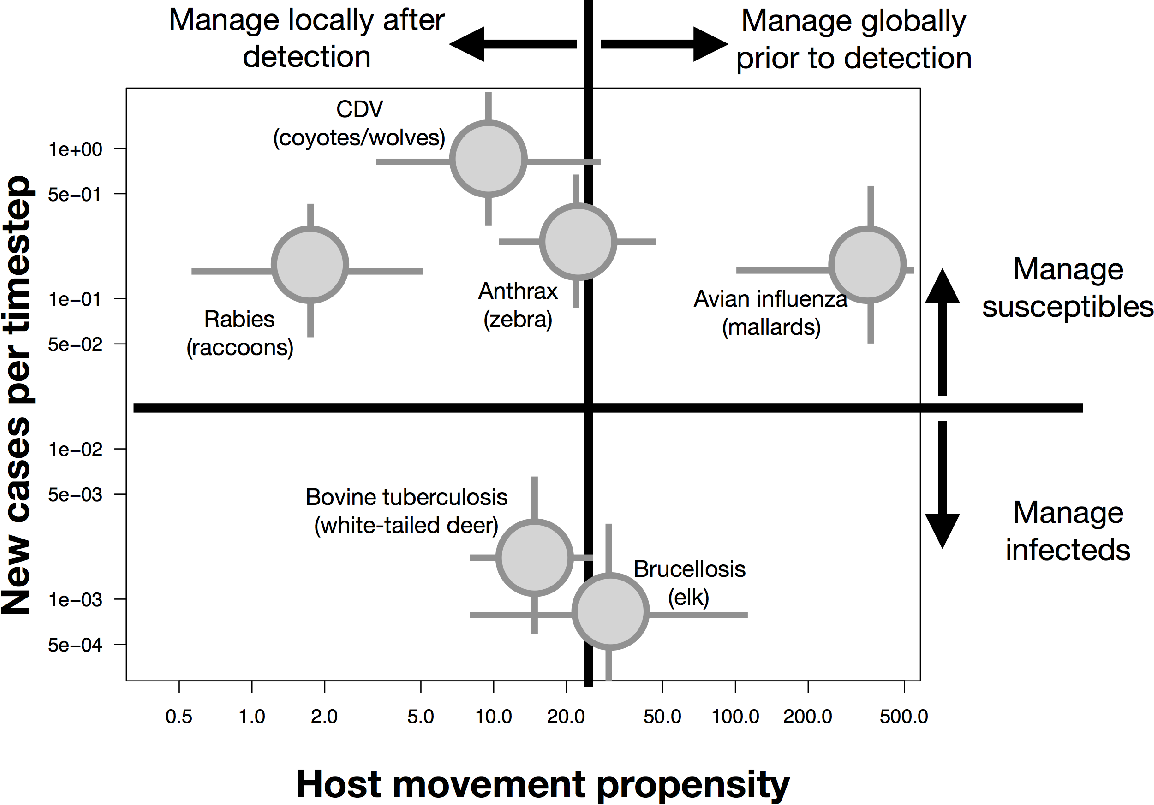
Hypothesised management efficacies across the disease propagation space. We partition disease propagation space along two axes: epidemic growth rate and movement propensity of the reservoir host (here, quantified as typical number of kilometers moved during the infectious period). Relative performance of various management strategies are indicated with arrows outside the illustration’s margins. Management actions that might fall into the upper-left quadrant include local depopulation of either host at infected premises. Management in the upper-right might include biosecurity measures targeting premises of either host, or prophylactic vaccination of the reservoir host. Management in the lower-right could include a wide range of actions. Many actions might perform well in the relatively easily-managed lower left-hand quadrant. System-specific values are indicated within the illustration. Values were left intentionally vague, and the values corresponding with the quadrant lines are arbitrarily chosen, since they will vary with ecological context, and with the particular organisms involved. Spatial extents indicated here are based on one important, current reservoir host listed below each point. The upper bound of movement for avian influenza has been curtailed. See Supplementary Materials: Table S1 for system-specific estimates.

Lastly, we expect that in some cases, many management actions may perform comparably. In these situations, relative economic and social costs of each action should factor heavily into management decision making.

Testing these conjectures requires us to explicitly incorporate epidemic growth rates and host movement propensities into a model of epidemic spread. Like many single-host models [6], our model considers management in the context of a dynamic disease transmission process, but we make that process spatially explicit, and allow new spillovers to emerge autonomously due to underlying infection dynamics in the reservoir host. Like many multi-host models [11, 12], magnitude of spillover risk is central to our model structure. However, where those models focus on the magnitude of risk in terms of force of infection from the reservoir host, we focus on that risk’s spatial extent. Spatially linking both hosts in one model allows us to compare a wider range of management actions that span both host species.

## 3 Spillover simulation model

Our pathogen propagation model includes three elements: transmission of the disease within species, transmission of the disease between species, and movement of species across space. The model operates on a 50 × 50 grid of spatial cells. We assume that host population sizes are fixed and identical in all occupied cells. The model uses a deterministic SIR disease process model with stochastic cell-to-cell host movements and between-species interactions to capture the effects of both host movement and epidemic growth.

Epidemic dynamics within a spatial cell follow the classic disease model of Kermack and McKendrick [14] without demography, which assumes that each individual’s disease status transitions from Susceptible (S) to Infectious (I) to Recovered (R). By keeping population sizes constant in all grid cells, we circumvent questions about whether within- and between-host-species transmission should be frequency or density-dependent. During an epidemic’s early exponential growth phase, the epidemic growth rate is equal to the difference between the transmission coefficient (*β*) and the recovery rate (*γ*) (Supplementary Materials: Section 1). We held recovery rate constant across all simulations, and systematically altered epidemic growth rates by manipulating *β* alone (Figure S1A).

The model’s spatial process starts with construction of an occupancy map for each host species, based on a stochastic selection of cells. The probability a cell is occupied by a particular host species is determined with a draw from a symmetric, bivariate Gaussian kernel centered on that host’s activity center (i.e., surrounding a spatial centroid), and with a variance equal to 50 (Figure 2A). If a previously drawn cell is chosen a second time for the same species, we redraw from the same distribution until a set of 1,000 unique cells is identified for each host (though both hosts can occupy the same cell). Spatial alignment between the host species is determined by the distance between host activity centers. This structure allows us to capture a gradient of spatial overlap between host species, from complete segregation to extensive spatial overlap throughout their ranges (Supplementary Materials: Section 4; Figure S2).

**Figure 2:**
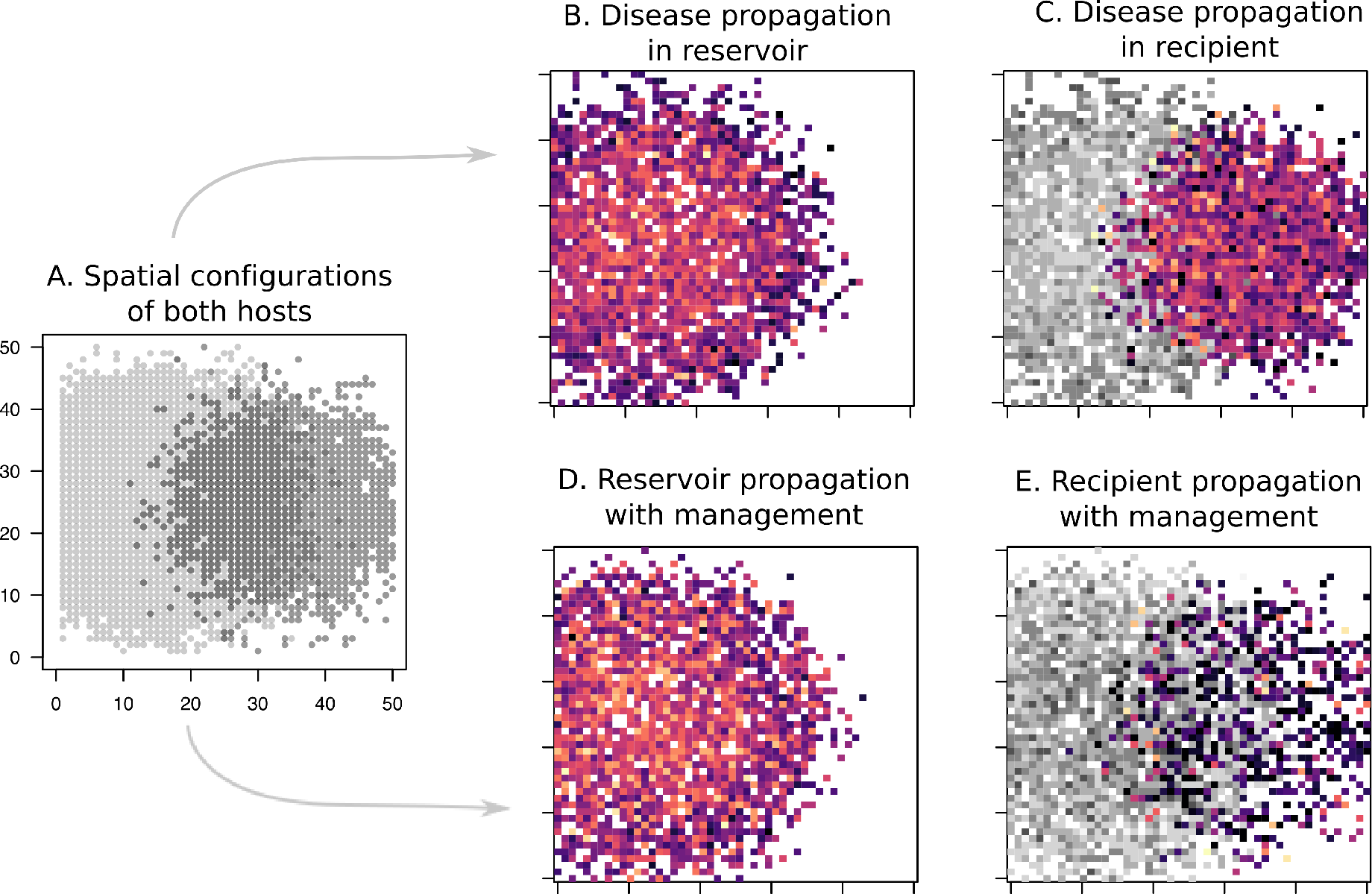
Simulation protocol. (A) Simulations begin by defining the spatial extent of the reservoir (light grey) and recipient (dark grey) hosts. A single infected individual is introduced into two reservoir host cells (B), with the structure of the subsequent epidemic determined by epidemic growth rate and host movement propensities. Here, lighter, more orange patches are infected earlier, whereas darker, more purple patches are infected later. C) The pathogen can then stochastically spill over to the recipient host in cells occupied by both host species (reservoir shown in grey; recipient in color), at a rate determined by the interspecific contact rate and the pathogen’s prevalence in the local reservoir host population. Management (here, retroactive vaccination of the recipient host) can alter the epidemic’s progression in both host species (D and E). Management actions are compared in terms of their ability to minimise the epidemic’s spatial extent in the recipient or reservoir host, and minimise the total number of recipient or reservoir host cases. Variation generated by manipulating spatial structure, disease propagation, and management efficacy are shown in Supplementary Figures S2, S3, and S4, respectively.

Individual reservoir and recipient hosts move stochastically between occupied grid cells at discrete time steps according to a tau-leap algorithm (Supplementary Materials: Section 2.1). We assume that these movements have a negligible effect on cell population densities, and hold population sizes constant within each occupied cell throughout the simulation. We only track the size of each disease compartment (i.e., proportion of susceptible, infected, and recovered individuals of each host species) within cells, not the disease status of particular individuals. Stochastic movement events by “individual” infected animals spark dynamics within newly-contacted cells, without altering cell densities.

Movement rates are normalised so that the same number of cell-to-cell movements is expected in all simulations, but the structure of these movements varies controllably. Movements are drawn from a monotonically decreasing function of Euclidean distance between cell centroids (i.e., a movement kernel, Supplementary Materials: Section 2.2), and we control host movement patterns by manipulating the distribution of movement distances (which we refer to as the host’s “movement propensity”). The probability that a dispersing host is infected is proportional to the local prevalence in that host’s population. Movement kernels with heavy tails correspond to high-dispersal systems (high movement propensity), while movement kernels with light tails correspond to low-dispersal (low movement propensity) systems (Figure S1B). Epidemics simulated under high movement propensities seed more new infections at a distance than epidemics arising under lower movement propensities [18], facilitating faster spatial spread. For simplicity, we apply the same dispersal kernel to both host species in all simulations presented here, and we do not allow colonisation of unoccupied cells. Arrival of a single infected host in an uninfected host cell always and instantly sparks a local epidemic in the newly contacted cell (Figure 2B).

Contacts between reservoir and recipient hosts are also treated as stochastic. The number of interspecific contacts is based on a pre-specified interspecific contact rate, which is held constant across all simulations. Contacts are then randomly assigned across all cells occupied by both reservoir and recipient hosts. The probability that an interspecific contact involves an infected reservoir host is proportional to the current infection prevalence in that cell’s reservoir host population. Though reservoir-to-recipient contacts are required to initiate epidemic dynamics in the recipient host population, we assume that recipient host populations experience no additional force of infection from reservoir hosts in local or neighbouring cells following the initial spillover event (Figure 2C). Stochastic movements between recipient hosts can then allow the epidemic to propagate throughout the entire recipient host population.

### 3.1 Management Actions

Our investigation focuses on six forms of management: prophylactic vaccination applied across an entire population regardless of current disease status, retroactive vaccination applied to infected cells and their neighbours following detection of disease, contact biosecurity, depopulation, spatial containment, and selective removal of infected reservoir hosts (Figure 2D, 2E). Details on implementation of all management actions within the model are contained in Table 1 and Supplementary Materials: Section 2.3. In all cases except for prophylactic vaccination, management is applied only to cells exceeding a specified threshold prevalence in the reservoir host (and to the direct neighbours of those cells for retroactive vaccination and depopulation; Table 1).

**Table 1:** Management structure within the model. *θ* is a user-controlled value indicating the prevalence at which management begins. **A** is the set of all cells with centroids within *ξ* units of the target cell (a set of "neighbour cells"; *ξ* was fixed to 3 for all simulations here). *N* is the number of reservoir hosts per occupied cell. *e*_*ij*_ is the movement rate between the *i*^*th*^ and *k*^*th*^ cells; *ι* is an alternative dispersal value applied to all cells outside the containment zone. It is fixed to 1/10,000 here.

| Action | Prevalence at which action begins | Effect of action on dynamical system | Spatial extent |
| --- | --- | --- | --- |
| Biosecurity | $\theta$ | $\beta I_{res} S_{recipient}$ becomes $\psi \beta I_{res} S_{recipient}$; $\psi \in [0, 1]$ | $\mathbf{A}$ |
| Prophylactic vaccination | N/A | $S$ reset to $S(1 - \nu)$ where $\nu \propto N_{cell}$ | All |
| Retroactive vaccination | $\theta$ | $S$ becomes $S(1 - \nu)$ where $\nu = 1 R_0$ | $\mathbf{A}$ |
| Test-cull | $\theta$ | $I$ becomes $\epsilon$ where $\epsilon \ll 1$ | $\mathbf{A}$ |
| Containment | $\theta$ | $e_{ij}$ becomes $\iota$ where $\iota \ll 1$ | $\mathbf{A}$ |
| Depopulation | $\theta$ | $I$ reset to 0 | $\mathbf{A}$ |

This management structure leaves ample room for development, including exploration of cost constraints and more complex management schemes. However, additional detail would require tailoring the model to a specific system, so here we simply present the overarching structure and leave further specification to future work.

### 3.2 Model process and outputs

We initiated the simulation by introducing the pathogen into two randomly chosen reservoir host cells at timestep 1, and we simulated the epidemic forward for 60 timesteps. We recorded the the time of first infection separately for reservoir and recipient hosts at each cell. Once a cell was infected, future pathogen introductions did not alter local disease dynamics. After the simulation, we calculated the proportion of reservoir and recipient cells that became infected, along with the maximum prevalence (or number of recovered cells, which could reflect disease-associated mortality burdens in some systems) over the full simulation. These metrics – spatial extent of the reservoir and recipient host epidemics, along with maximum epidemic size, and total disease-induced mortalities in both hosts – provided a basis for comparing disease propagation dynamics under varying rates of host movement and epidemic growth. The model was implemented as a *de novo* simulation in R [15].

## 4 Identifying the most effective management strategies

In order to identify which management action produced the best results under particular conditions of host movement propensities and epidemic growth, we ran simulations according to a full-factorial design (Supplementary Materials: Section 3; Table S1). Parameters varied along the following six dimensions: 1) epidemic growth rate; 2) variance and kurtosis of the hosts’ dispersal function (“movement propensity”); 3) prevalence at which management began; 4) host density within each cell; 5) distance between reservoir and recipient host activity centers; and 6) management objective of interest. The objectives we considered were minimising spatial extent of the epidemic in the recipient host; minimising total number of recipient host cases; minimising spatial extent of the epidemic in the reservoir host; or minimising total number of reservoir host cases. We focus primarily on spillover management in the recipient host population, but include results from objectives 3 and 4 in Supplementary Materials: Section 5.2.

We first explored raw output values over the disease propagation space (Figure 3), and then tabulated which management action performed “best” (i.e., minimised epidemic size or epidemic spatial extent in the reservoir or recipient host) at every parameter combination (Figure 4A-4D). We then used logistic regression to correlate when a management action was ranked as the best (yes or no) with epidemic growth rate and tail weight in the host’s dispersal kernel. All simulations contributed to model fits, so that the full-factorial design balanced inferences over a range of values for reservoir population density, spatial divide between host activity centers, and prevalence at which management began.

**Figure 3:**
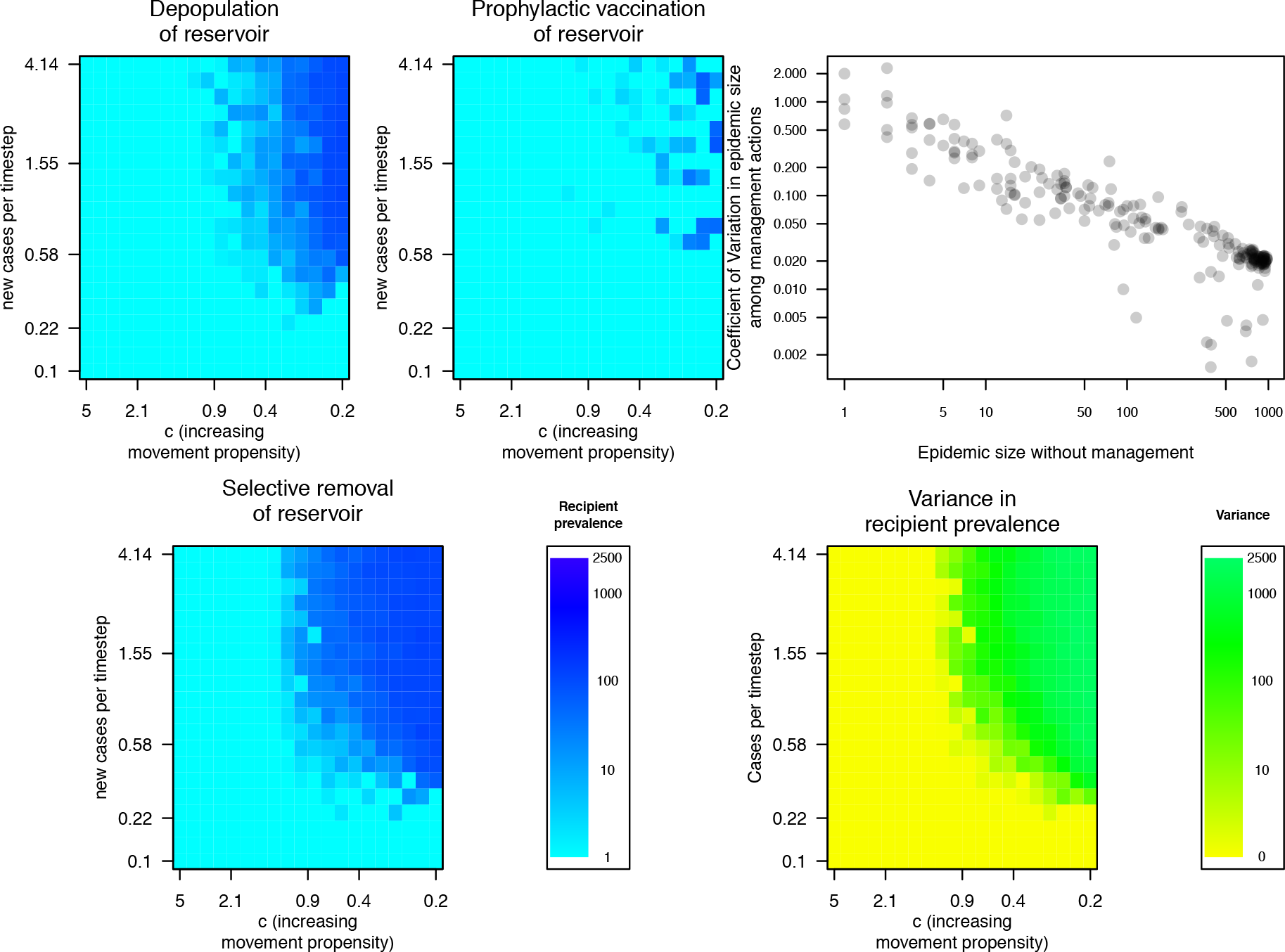
Simulation output. Heat maps (A-C) show the aggregate recipient host prevalence under three different management actions along the dimensions of epidemic growth rate (Cases per timestep) and movement distance (movement propensity increases to the right): (a) depopulation of the reservoir host; (b) prophylactic vaccination of the reservoir host; and (c) selective removal of the reservoir host. In cases where unmanaged epidemics were large, outcomes varied dramatically among management actions, and a clear “best action” was identifiable (this is the case among the three sets of actions shown here, in which prophylactic vaccination of the reservoir clearly outperforms the other two actions for the fastest-growing, fastest-spreading epidemics). However, in cases where unmanaged epidemics were small, many actions performed comparably (d). (e) Variance between management outcomes increased with both movement and the number of cases per timestep. Simulations in panels A-C partition both epidemic growth rate and c into 20 blocks, fix management initiation prevalence to 0.01, and set the spatial divide between host centroids to 30 cells.

**Figure 4:**
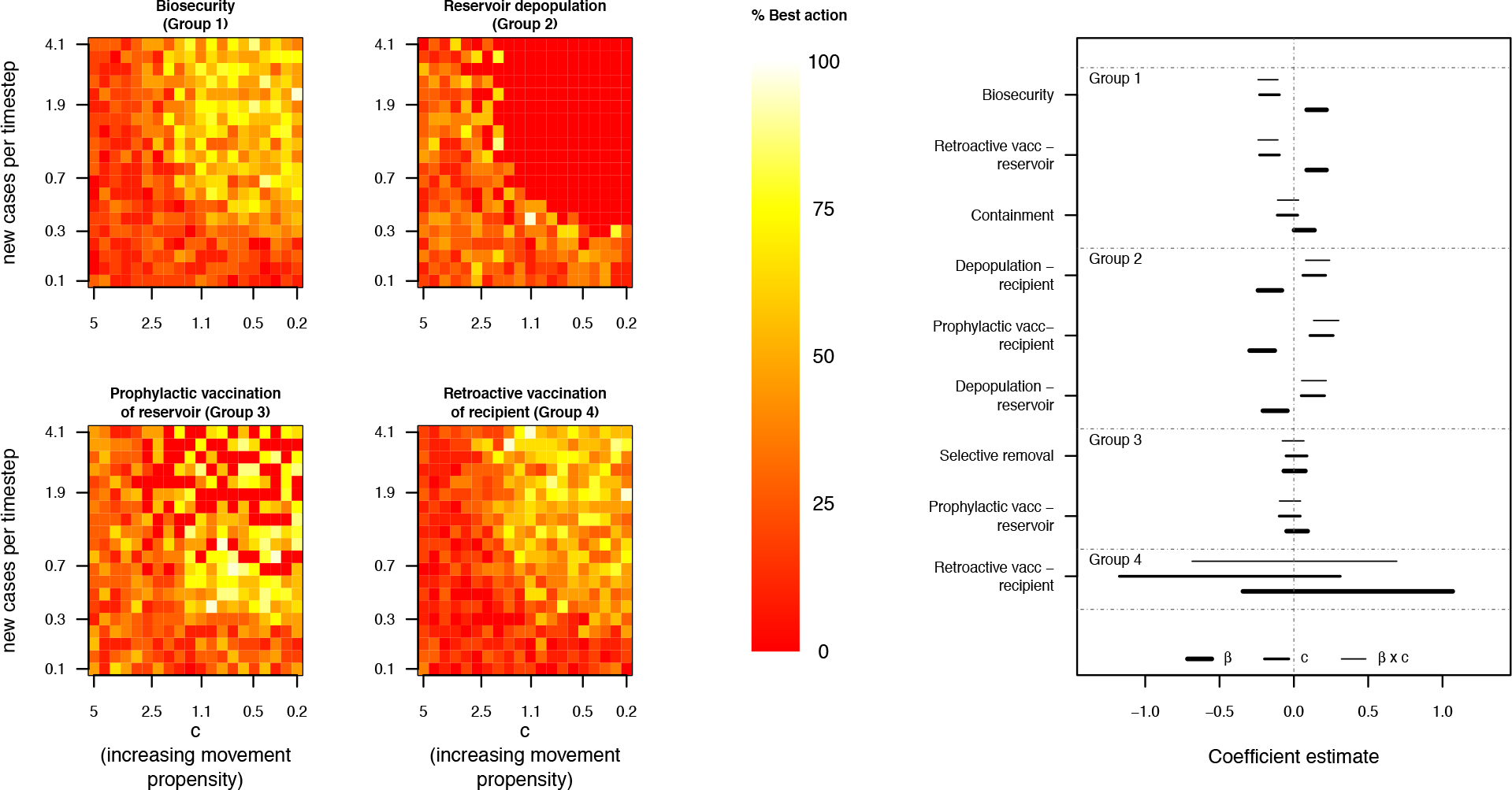
Comparison of management actions over the disease propagation space. The left four panels show regions of the parameter space where four representative actions performed well at controlling spillover. The output metric over which actions are compared here is the number of recipient host patches infected over the entire simulation. Parallel figures for the other three outcomes are shown in Figures S6-S8. Pixel colors represent the simulated, raw proportion of times that a given action performed better than all other actions at that particular combination of epidemic growth rate and movement distance. Lighter, more yellow regions represent areas where the action was most likely to perform best, whereas red regions represent parameter values where the focal action was consistently outcompeted by other options. The right panel shows coefficients linking epidemic growth rate and movement propensity to the probability that each management action (color-coded) performs best under each output metric. Lines extend to 95% confidence intervals around each coefficient’s point estimate. All fits were generated from models that also included a term for spatial divide between reservoir and recipient host activity centers, but we omitted that term in the figure for simplicity. Complete model results are included in the Supplementary Materials: Section 5.2. Simulations in panels A-D partition both epidemic growth rate and c into 20 blocks, fix management initiation prevalence to 0.01, and set the spatial divide between host centroids to 30 cells.

Next, we grouped management actions according to their coefficient estimates from the logistic regression (Figure 4), and compared conditions of strong relatively performance for each group to our expectations in Figure 1.

## 5 Results

Our simulator produced a wide range of spatiotemporal spillover and transmission dynamics (Figure 3; Figures S3-S4), and these dynamics generally responded as expected to the various management interventions (Figure 3; Figure S5). While the results presented here focus on one objective, limiting the total number of infected cells in the recipient host population, similar patterns were observed for limiting the total number of recipient hosts infected (Supplementary Materials: Section 5.2). Unsurprisingly, we saw some deviations from these patterns when objectives were centered on the reservoir host. For instance, we allowed biosecurity to only target interspecific contacts, so it had no bearing on disease dynamics in the reservoir host. Most of the observed deviations between the reservoir and recipient host objectives could be readily explained with similar logic.

Action performances typically fell into two groups when aggregated across the entire parameter space, but which actions fell into which group depended on objective (Table S3). When the objective was minimising the epidemic’s spatial extent in the recipient host population, the better-performing group consisted of biosecurity, prophylactic vaccination of the reservoir host, and retroactive vaccination of either host (Table S3). Epidemics tended to be easily controlled by many management strategies when movements were mostly local and epidemic growth rates were low (Figure 3, see Supplementary Materials: Section 5.1 and 5.2 for statistical results, and Supplementary Materials: Section 5.3 for results under other objectives).

Actions could be grouped according to how their relative performance related to epidemic growth rate and host movement propensities. Biosecurity, containment, and retroactive vaccination of the local reservoir host (Group 1), which all aim to limit spillover effects by controlling interspecific contact or spatial propagation, performed best for fast-growing epidemics in hosts with high movement propensities (the upper right-hand corner of Figure 1). Depopulation of both hosts, along with prophylactic vaccination of the recipient host (Group 2) had the strongest relative performance in scenarios where epidemics grew quickly, but host movement propensities were low. The relative performance of selective removal and prophylactic vaccination of the reservoir (Group 3), both of which target prevalence and epidemic growth rates in the reservoir population, did not structure with epidemic growth rate or host movement propensities. Management groupings changed with alterations in the objective function, and this was particularly true when the measured objective focused on a different host. For instance, when the objective was to minimise infected reservoir patches, selective removal performed best when epidemics grew slowly, whereas prophylactic vaccination of the reservoir was best when epidemic growth rates were high, even though these actions were grouped together when the objective was to minimise epidemic size ore extent recipient host (Figure S7).

## 6 Discussion

Pathogen spillover at the wildlife-livestock interface is a persistent and expensive problem for food security and wildlife conservation alike. While pathogen spillover dynamics have been explicitly studied in many wildlife-livestock systems, disease ecologists lack a general framework for considering the context in which each management action should perform best. Here, we presented a first effort toward modeling spatially explicit spillover management across both reservoir and recipient host species.

Our simulation results usually agreed with our *a priori* intuition about how management would interact with spatially explicit epidemic propagation. Epidemic growth rate and host movement propensities interacted to generate variation in epidemic size and spatial extent (Figure 4; Figure S6-S8), and epidemic growth rate was generally the most powerful determinant of which management action performed “best” (Figure 4E; see Supplementary Materials: Section 5.3 for additional results). While our findings are likely sensitive to the range of parameter values explored (epidemics sometimes failed to propagate in one or both hosts in a substantial region of the parameter space), they were nevertheless consistent with the basic categorisation proposed in Figure 1. In fast-growing, high-movement-propensity systems, limiting and localising recipient host interactions with infected reservoir hosts — whether by reducing reservoir prevalence through vaccination, or by reducing interspecific contact rates through biosecurity — was the best strategy for recipient-focused objectives. Containment also performed relatively well in these scenarios, except in cases where host movements consistently exceeded the size of the containment region (though the implications of varying containment region size was not explored here). Actions like depopulation or prophylactic vaccination of the recipient host, which limit epidemic growth by controlling the size of the susceptible pool without targeting movement, performed relatively better when epidemic growth rates were high and movement propensities were low. These actions might be favored for fast-growing, low-movement-propensity epidemics, and may be competitive in higher-movement-propensity scenarios, depending on cost. Performance of actions that controlled local prevalence in reservoir host populations through prophylactic vaccination or selective removal did not exhibit strong patterns with epidemic growth rate or host movement propensities. However, retroactive vaccination of the reservoir host performed comparably in that same ecological scenarios, and may be more politically and socially palatable when feasible (Table S3; **Sokolow et al. *in revision***).

All management actions were competitive with one another when epidemic growth rates were slow, clearing a path for cost to play a larger role in decision-making (Figure 3D, 3E). This partially held for fast-growing epidemics with low host movement propensities as well, and has been demonstrated in detail in several wildlife-livestock spillover diseases. For instance, management models of foot-and-mouth disease (which we might categorize as a fast-growing disease with a lower movement propensity) have investigated a wide range of different management strategies and found that a variety of actions might be deemed appropriate, depending on the specific objective, the action’s cost, and the prevalence at which management begins [16, 17].

We were somewhat surprised that dispersal kernel structure did not play a more powerful role in shaping management efficacy here (Figure 4E). This could be because we held the total number of cell-to-cell movements constant throughout the simulations. In reality, host species vary in both the structure and the number of moves they make. We anticipate that movement may thus play a more powerful role than these results suggest, but this question merits additional follow-up.

We elected to emphasize epidemic growth rate instead of the pathogen’s basic reproductive ratio (*R*_0_) because disease management depends on calendar time rather than the pathogen’s generational timespan. Management requiring construction of fencing for biosecurity, or depopulation of infected premises, would be much more effective in a system with a slow epidemic growth rate than a rapid one, even if the epidemics were identical in terms of *R*_0_. This could play out empirically, for example, in bovine tuberculosis (bTB) management in Michigan, USA, where models suggest that a relatively constant, but low level of spillover pressure from wildlife could be successfully mitigated through fencing [18]. It is the slow growth rate, and not that the basic reproductive ratio, that renders fencing feasible for Michigan bTB. Similarly slow measures would be less effective for a pathogen with a comparable *R*_0_(estimated to be around 1.5 for Michigan bTB), but a shorter infectious period. Contrast bTB management, for instance, with plausible management for the 2014 Ebola outbreak, which also exhibited a basic reproductive ratio around 1.5, [19].

### 6.1 Framework limitations

This model was designed around the assumptions most relevant to our particular question of interest, namely, how allocation of disease management effort should vary along a two-dimensional continuum of epidemic growth rate and host movement propensity. Many other facets of the pathogen spillover and management decision-making process were simplified to isolate this question. In particular, the disease process model is subject to the same constraints facing many SIR models: we assumed that immunity is lifelong; that disease does not induce mortality; that host densities are constant through space and time; and that disease-related rates like recovery and transmission are effectively constant across all individuals. Additionally, our selection of timescale and epidemic duration were arbitrary. We assumed that host movements were random and independent, and that both host species moved according to the same movement kernels. All of these assumptions are unrealistic in some scenarios, and we discuss each in greater depth in the Supplementary Materials: Section 6.

Additionally, our results are driven in part by the assumed efficacy of each management action. Efficacy values were generated from a set of preliminary simulations identifying parameter values that generated similar effect sizes. A full sensitivity analysis incorporating all model parameters is beyond our current scope. However, we recognize that such a study is a critical next-step. A simplified tactic, in which we sample randomly over an ungridded (but uniform) parameter space to assess output sensitivities of the various parameters, is the subject of current investigation. At this time, we simply emphasize that our current objectives are also fundamentally different then those of a researcher aiming to forecast management efficacy in a given system. To compare management for specific systems where the general process parameter space is already quite constrained, a more detailed investigation of sensitivities to cost, efficacy, etc. would be important.

We only allow for a single management action to be undertaken at a time, and that we do not account for costs or logistical constraints associated with those actions. Costs vary dramatically across systems and contexts (see, for example, the discussion in [20] surrounding the costs of brucellosis management), and placing any specific cost on an action quickly constrains the set of systems to which the results extend. At the same time, cost-benefit trade-offs are already being used to justify spillover disease management in reservoir hosts. For example, Sterner et al. [21] argue that even though oral rabies vaccination in the U.S. and Canada are quite costly, those aggregate costs are lower than the costs of post-exposure prophylaxis that would be required to manage rabies in the spillover host (here, humans). Further comparative inquiry into cost is increasingly necessary.

Beyond economic costs, disease management logistics take time to coordinate, and this constrains how quickly management follows detection, and how many cells can be managed at once. From a political stance, reservoir and recipient host management decisions are often determined by separate agencies with distinct, and sometimes discordant, objectives. Being able to weigh disease management actions against more complex objective functions that include those different viewpoints is an important extension that would allow managers, researchers, livestock producers, and conservationists to reach some common ground.

Finally, our model does not allow management actions to fundamentally alter the system’s underlying ecology (i.e., to pull “ecological levers”, **Sokolow et al. *in revision***). In reality, some of these actions — particularly, depopulation and selective removal — have the potential to impose major and lasting alterations. Allowing management to perturb underlying system ecology is an important issue for future exploration.

### 6.2 Framework extensions

The model’s simple structure means it can generate initial expectations about epidemic progression in understudied systems, even before the specific epidemiological process is well-understood. Accounting for disease and movement dynamics in both reservoir and recipient hosts allows us to compare a broader suite of management actions available for constraining emerging disease. Rapid parameterisation of the model may often be feasible, since both epidemic growth rate and host movement structure can be inferred from data available shortly after an epidemic’s inception. As system-specific information accumulates, the modeling structure could be refined to encapsulate emerging detail about how, when, and where to optimally manage the disease [22]**Carpenter et al. 2011 Vet Diag Invest**.

If multiple management options are available, coupling actions that tackle different aspects of transmission (e.g., actions from contrasting groups in Figure 4) may be beneficial. For instance, reservoir-focused actions can reduce the overall spillover risk by limiting the spatial extent and prevalence of the pathogen in the reservoir. However, unless these actions completely eliminate spillover risk, they may be most effective when coupled with targeted, responsive management in the recipient host (e.g., depopulation or retroactive vaccination). This both-hosts approach has already been used to manage several slow-growing, low-movement-propensity pathogens at the wildlife-livestock interface (e.g., brucellosis management around the Greater Yellowstone Ecosystem **Scurlock**, or bovine tuberculosis management in Minnesota, USA white-tailed deer [23]).

Lastly, the best-performing management in any context also depended on the objective. This is consistent with a rich body of work on adaptive management and structured decision making in a range of ecological contexts **[27]** including disease dynamics [17]**[28]**. In reality, objectives likely differ for the two host species, with recipient management focusing around limiting aggregate case load, and reservoir management emphasizing detection and mitigation of prevalence pulses(**Plowright et al. *in revision***). Given the extensive discussion of objective functions elsewhere, however, we do not dwell on them further here.

### 6.3 Conclusion

The risk and burden of pathogen spillover depends on a spatially explicit disease propagation process that operates in both reservoir and recipient host populations. Our model provides a useful starting point for planning management of disease at the wildlife-livestock interface based on general epidemiological traits of the system. It underscores the critical role that epidemic growth rate and spatial context play in determining management efficacy, and could be tailored to the specifics of a wide variety of pathogen spillover systems.

## Additional Information

### Data Accessibility

All estimates included in Figure 1 are tabled in the Supplementary files, and simulation and analysis code is contained in the supplement.

### Authors’ Contributions

KRM and PCC developed the conceptual framework, and all authors contributed to model refinements. KRM and LSM constructed the simulation model, and KRM, LSM, and PCC developed the model analysis. KRM drafted the manuscript, and all authors contributed to multiple rounds of manuscript revision.

### Competing Interests

We have no competing interests.

### Funding

LS was supported by the L’Oreal For Women in Science Fellowship. KMP was supported by USDA.

## Acknowledgements

Any use of trade, firm, or product names is for descriptive purposes only and does not imply endorsement by the U.S. Government.

