## Supplementary Materials for "Epidemic Growth Rates and Host Movement Patterns Shape Management Performance for Pathogen Spillover at The Wildlife-livestock Interface"

April, 2019

**3 Parameters and parameter space explored** **7**

**4 Assessing model performance** **8**

**5 Additional results** **12**

**References** **25**

*Disclaimer: This draft manuscript is distributed solely for the purposes of scientific peer review. Its content is deliberative and predecisional, so it must not be disclosed or released by reviewers. Because the manuscript has not yet been approved for publication by the U.S. Geological Survey (USGS), it does not represent any official USGS finding or policy.*

### 1 Empirical estimates of epidemiological rates

| **Disease** | **Exponential growth rate** | $\beta$ | $\gamma$ | **Potential movement while infected (km)** |
|---|---|---|---|---|
| Brucellosis | 0.0006 [1] | NA | NA | 3-8 km/y [2] |
| Bovine Tuberculosis | 0.002 ($R_0 \approx 2.59$ [3]) | NA | NA | 15 6km/y [4] |
| Rabies | 0.152; $R_0 \approx$ 2-2.44 [3, 5] | 0.18 | 35 | 3km [6] |
| Avian influenza | 0.18; $R_0 \approx 2.24$ [7, 8] | 0.0078 [9] | 7d [10] | 584-712 km [11] |
| Canine distemper (CDV) | 0.42 | 0.16-0.30 [12] | 15-23d acute; 67-74d persistent [12, 13] | 17.3 km (based on a mean pack range size of 300km$^2$) [14] |
| Anthrax | 0.46; $R_0 \approx$ 2.98-5.97 [15] | 22.5 | 7.5 d [16] | 3 km/d |

Table S1: Estimated epidemic growth rates and host movement potentials for systems shown in Figure 1 of the main text. Undoubtedly, an SIR process model is insufficient for any one of these systems. However, SIR-based estimates may still be sufficient for the coarse classification we aim to make on the simple basis of epidemic growth rate and host movement.

### 2 Model structure

The model begins with a spatial grid of 50x50 cells that could approximate counties in the U.S. (we envision 30mi x 30mi grid cells, but the exact spatial extent of the cells does not affect the simulations). The disease process within each cell follows the model of Kermack and McKendrick (1927). This model rests on a set of three ordinary differential equations (ODEs) describing how individuals within a population move from Susceptible (S) to Infected (I) to recovered

(R) states.

$$\frac{dS}{dt} = -\beta SI \tag{1}$$

$$\frac{dI}{dt} = \beta SI - \gamma I \tag{2}$$

$$\frac{dR}{dt} = \gamma I \tag{3}$$

The epidemic growth rate can be determined by using the Jacobian of the set of ODEs, solved at state values corresponding to the disease-free equilibrium. The Jacobian of the ODEs is simply the derivative of each equation in the set with respect to each variable. Here,

$$J = \begin{bmatrix} -\beta I^* & -\beta S^* & 0 \\ \beta I^* & \beta S^* - \gamma & 0 \\ 0 & \gamma & 0 \end{bmatrix} \tag{4}$$

Thus,

$$\det(\mathbf{J} - \mathbf{\Lambda I}) = \begin{vmatrix} -\beta I^* - \Lambda & -\beta S^* & 0 \\ \beta I^* & \beta S^* - \gamma - \Lambda & 0 \\ 0 & \gamma & 0 - \Lambda \end{vmatrix} \tag{5}$$

$$= (-\beta I^* - \Lambda) \begin{vmatrix} \beta S^* - \gamma - \Lambda & 0 \\ \gamma & 0 - \Lambda \end{vmatrix} - (-\beta S^*) \begin{vmatrix} \beta I^* & 0 \\ 0 & 0 - \Lambda \end{vmatrix} + 0 \begin{vmatrix} \beta I^* & \beta S^* - \gamma - \Lambda \\ 0 & \gamma \end{vmatrix} \tag{6}$$

$$= (-\beta I^* - \Lambda)(\beta S^* - \gamma - \Lambda)(-\Lambda) + \beta S^* \beta I^* (-\Lambda) \tag{7}$$

Substituting in the $S^*$ and $I^*$ values for the disease-free equilibrium, $S^* = 1$ and $I^* = 0$, leaves:

$$(-\beta \times 0 - \Lambda)(\beta \times 1 - \gamma - \Lambda) + \beta(1)\beta(0)(-\Lambda) \tag{8}$$

$$= -\Lambda(\beta - \gamma - \Lambda) \tag{9}$$

Roots occur at $\Lambda = 0$ and $\Lambda = \beta - \gamma$. The latter solution is the epidemic growth rate.

#### 2.1 Tau-leap implementation of stochastic movement

We use a tau-leap algorithm to simulate cell-to-cell movement by reservoir hosts. The tau-leap is a two-step approximation that extends the Gillespie algorithm to operate on discrete and systematic units of time, $\Delta t$, while also allowing

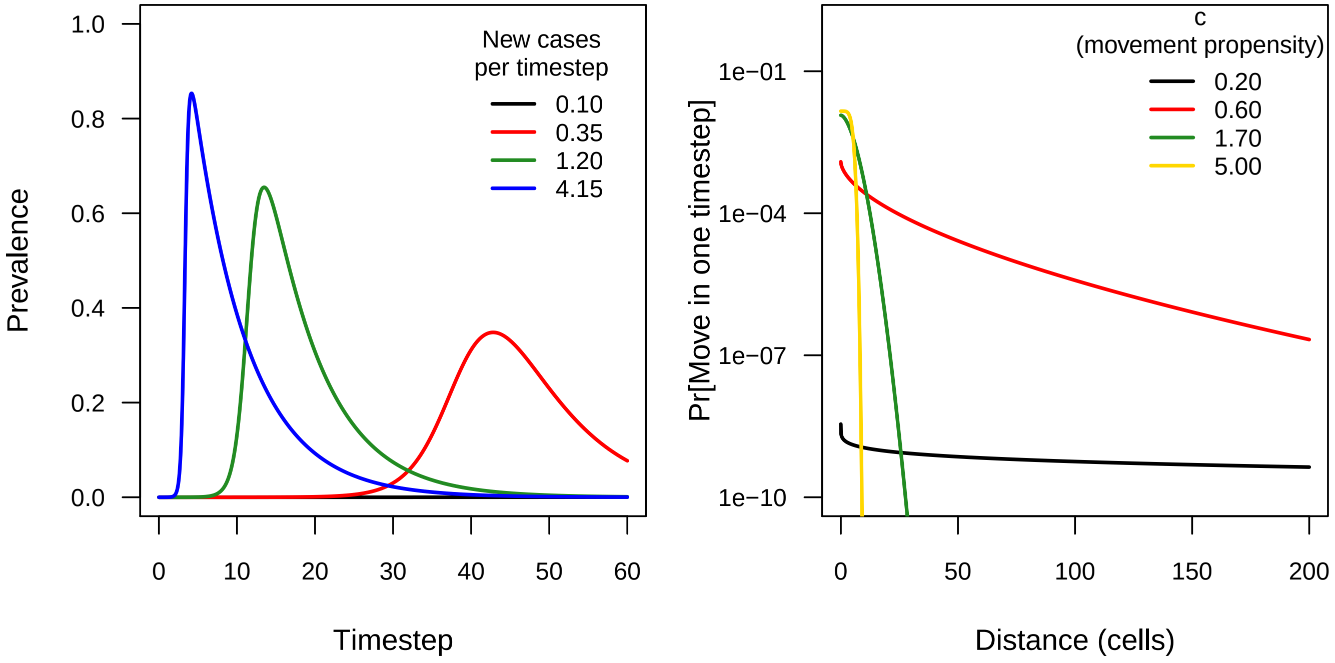

Figure S1: A) Disease dynamics under the range of epidemic growth rates explored here. B) Dispersal kernels under the range of $c$-values explored here. Note that the tailweight declines as $c$ increases.

updates to occur in blocks (this is a useful feature in our scenario, since we are interested in a timescale where waiting times between moves are quite low, such that simulating waits between specific moves is computationally impractical). The first step of the tau-leap is to draw the number of moves during $\Delta t$, $N_{\text{events}}(\Delta t)$, from a Poisson distribution, such that

$$N_{\text{events}}(\Delta t) \sim \text{Poisson}(\lambda), \tag{10}$$

$$\lambda = \sum_{i,j} \text{move}_{ij} \tag{11}$$

The second step is to then draw the originating and terminal cells associated with each move, Connect$(\Delta t)$, generated by a draw of size $N_{\text{events}}(t)$ from a multinomial distribution with probabilities equal to $p_{ij}$.

$$\text{Connect}_1(\Delta t) \sim \text{Multinom}(N_{\text{events}}(\Delta t), p_{ij}) \tag{12}$$

(13)

Because this is a multinomial draw, several moves between the same pair of cells can occur within a single unit of time; the frequency of moves remains in proportion to the pairwise distance between cells. We assume that $p_{ij} = p_{ji}$, though this would not necessarily be the case. Thus, after drawing the pair of connected cells, we then assign a direction to all moves using a single of draw of size $N_{\text{events}}(\Delta t)$, from a binomial distribution with success probability of .5.

$$\text{Direction}(\Delta t) \sim \text{Binomial}\left(N_{\text{events}}(\Delta t), .5\right), \tag{14}$$

$$\text{Connect}(\Delta t)[i] = \begin{cases} \text{Connect}_1(\Delta t)[i] & \text{if Direction}(\Delta t)[i] = 0, \\ \text{rev}(\text{Connect}_1(\Delta t)) & \text{otherwise} \end{cases} \tag{15}$$

Once the originating cells are determined, we then simulate the infection status of the moving host, which is determined by a Bernoulli trial associated with each move in Connect($\Delta t$), with success probabilities defined by the current prevalence (number of infectious individuals) present in each connection's originating cell. Prevalence is in turn determined by identifying the current "infection age" of the originating cell (that is, the difference between current time and time at which the cell was first infected), and solving for $I$ in the equations 1-3 at that infection age.

This progression of events produces a stochastically determined set of moves, with every move assigned a corresponding binary infection status (infectious or not). Any movement of an infectious host initiates the deterministic disease transmission process in the terminal cell. Throughout the simulation, we assume cell-specific population sizes remain constant, and we assume that cell-to-cell movement probabilities do not change throughout the simulation.

#### 2.2 Tail weight in the dispersal function

In order to modulate the movement distribution, we used a family of dispersal kernel distributions that could range from exponential to leptokurtic (i.e., fat-tailed [17,18]). This flexibility is important, since well-characterized dispersal kernels for animal species vary, but are often heavy-tailed (e.g., [19, 20]). Let $f\left(\text{dist}_{ij}\right)$ be the dispersal probability between two points at a fixed distance $\text{dist}_{ij}$. Then

$$f\left(\text{dist}_{ij}\right) = \frac{1}{N}\exp\left[-\left(\frac{\text{dist}_{ij}}{\alpha}\right)^c\right]$$

In this formulation, $\alpha$ is a parameter describing the dispersal distance, $c$ is a shape parameter controlling the distribution's kurtosis, and $N$ is a normalization constant that can be written as

$$N = \frac{2\pi\alpha^2\Gamma\left(\dfrac{2}{c}\right)}{c}$$

where $\Gamma(x)$ is the usual $\Gamma$ function. For $c \leq 1$, the distribution is fat-tailed; it is exponential at $c = 1$, and platykurtic at $c > 1$ (for reference, the Gaussian distribution has c = 2). Cell-to-cell movement probabilities, $p_{ij}$, are proportional to $\text{dist}_{ij}$, normalized across all potential moves. We rescale the weights of all distributions so that the total number of expected moves is held constant across all values of c. Dispersal is taken to be symmetric in all directions.

#### 2.3 Implementation of management actions

##### 2.3.1 Prophylactic vaccination

Both forms of vaccination work primarily on the deterministic side of the disease transmission model, by effectively lowering the disease's $R_0$ value ($p_c = 1 - \frac{1}{R_0}$) [3]. Prophylactic vaccination operates prior to spillover, and alters the proportion of susceptible hosts. We simulate prophylactic vaccination by shifting the initial conditions for the transmission process, so that some proportion of the host population starts in the "recovered", as opposed to "susceptible", state. Prophylactic vaccination is applied at a constant rate (equal to the proportion require to achieve herd immunity based on the system's $R_0$) across all occupied host cells over the entire spatial domain. This strategy is being used in an effort to manage spillover risk for avian influenza and rabies. We explore its implications when applied to either the reservoir or the recipient host species.

##### 2.3.2 Retroactive vaccination

Retroactive vaccination consists of responsively vaccinating hosts in reaction to pathogen detection. In our simulation, this is mathematically identical to prophylactic vaccination (some proportion of the susceptible recipients are shifted to the recovered category), but instead of vaccinating all premises to the same level, we vaccinate only the patches in which the infection itself reaches a predetermined threshold, and those patches' direct neighbors. Vaccination is applied at the proportion required to achieve herd immunity, as determined by $R_0$. Retroactive vaccination can be applied to either the reservoir or the recipient host. Retroactive vaccination does not completely eliminate the pathogen.

##### 2.3.3 Contact biosecurity

Contact biosecurity consists of actions like improving fencing and removing attractants, and aims to reduce the rate of direct contacts between the reservoir and the recipient hosts in the same patch. We simulate contact biosecurity actions by reducing the probability of interspecific contacts by a fixed constant in patches where "biosecurity" is applied. As with retroactive vaccination, biosecurity is applied when pathogen prevalence crosses a pre-specified threshold in the reservoir host. We take biosecurity to reduce the interspecific contact rate by 90% in all simulations here.

##### 2.3.4 Selective removal

selective removal strategies alter disease transmission by reducing prevalence in the reservoir host population. This strategy has been tested, for example, to manage brucellosis in elk in parts of Wyoming (Scurlock report), and to improve bighorn sheep population growth rates following disease spillover events. We simulate selective removal by reducing reservoir host prevalence in targeted patches by a particular amount - that is, the proportion of reservoir hosts in the infected category is lowered. This affects the deterministic disease dynamics. Isolation and removal of symptomatic individuals is a special case of selective removal, with no testing cost. Here, we only apply selective removal to the reservoir host.

###### 2.3.5 Depopulation

Depopulation consists of complete removal of all animals of the specified host species from a given cell and its neighbors within the management radius. Depopulation is followed by instantaneous restocking of all depopulated cells with susceptible hosts, so it has no effect on cell density. This is a reasonable assumption if spillover events do not lead to massive, species-wide reductions in host densities.

###### 2.3.6 Containment

Under the containment strategy, we first identify all cells within a fixed distance of the epidemic's starting point. Since disease is deterministic in this model, cells where the epidemic begins will always cross the prevalence detection threshold first, so the initiating cells are the same as the cells where infection is first detected. We then group the cells into a community of cells within the "containment" zone, and a community of cells outside that zone. We completely eliminated all moves between the two groups for both the recipient and the reservoir host populations.

#### 3 Parameters and parameter space explored

The simulator has 17 parameters, six of which we varied systematically in our simulation study. A complete list of all parameters is included in Table S2. We systematically varied the six investigated parameters in a full-factorial design, and ran a replicate simulation at each combination.

| Parameter | Description | # levels | Values investigated |
|---|---|---|---|
| $\beta$ | Determines epidemic growth rate (expected # new cases per day) | 4 | 0.10, 0.34, 1.20, 4.14 |
| c | Dispersion parameter of movement kernel | 4 | 0.20, 0.58, 1.71, 5.00 |
| $\gamma$ | Recovery rate | 1 | 1/7 |
| Management | Management action implemented | 9 | Depopulation of the reservoir<br>Depopulation of the recipient<br>Prophylactic vaccination of the reservoir<br>Prophylactic vaccination of the recipient<br>Biosecurity<br>Selective removal<br>Retroactive vaccination of the reservoir<br>Retroactive vaccination of the recipient<br>Containment<br>None |
| $N$ | Size of the reservoir host population within each cell | 4 | 10, 368, 13572, 500,000 |
| $\tau_{\max}$ | Number of timesteps simulation ran | 1 | 60 |
| $X_{\text{in}}$ and $Y_{\text{in}}$ | $X-$ and $Y-$ dimensions of the grid defining the simulator's spatial extent | 1 | 50 |
| $\xi$ | Spatial radius defining the neighborhood of cells **A** within which management is applied | 1 | 3 |
| $\psi$ | Biosecurity efficacy from Table S2 above | 1 | 0.1 |
| $\nu$ | Proportion of individuals in a cell who receive prophylactic vaccination | 1 | $1 - \frac{1}{R_0}$ |
| $\theta$ | Prevalence that initiates management | 4 | 0.001, 0.01, 0.10, 1.00 |
| Spatial divide | Distance (number of cells) between population centroids of reservoir and recipient host populations | 4 | 0, 15, 30, 45 |
| $\iota$ | Containment distance | 1 | 1/10,000 |
| | Cross-species contact rate | 1 | Within-species contact rate between cells 2 units apart according to the specified movement kernel |
| | Premise size (number of animals of the recipient host species per occupied cell) | 1 | $N$ |
| | Multiplicative constant adjusting the weighting in the movement kernel to generate an appropriate number of moves | 1 | 10,000 |
| $\alpha$ | Term structuring the dispersal kernel | 1 | 5 |

Table S2: Parameters that were systematically varied during the simulations, along with values investigated. Simulations presented in main text figures were run on a finer grid of twenty partitions each along $c$ and epidemic growth rate.

### 4 Assessing model performance

We assessed model performance visually under a wide range of parameter combinations to be sure the simulator performed as expected. We first evaluated whether our spatial configuration protocol worked properly by plotting occupancy patterns for the reservoir and recipient host under varying levels of spatial divide (Figure S2).

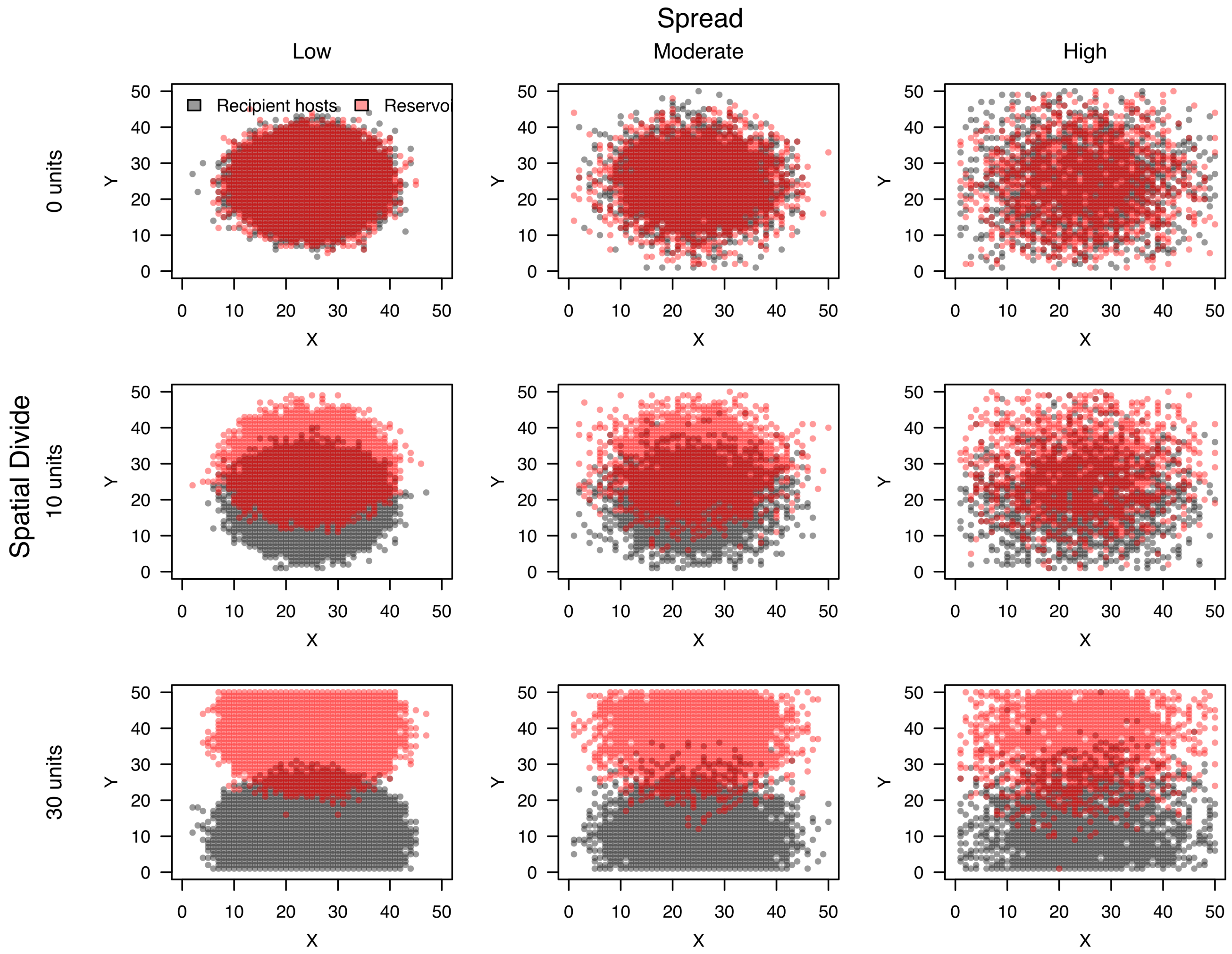

Figure S2: Spatial configurations of reservoir and recipient hosts under different values of “spread” and “spatial divide”.

We then plotted reservoir and recipient host epidemic dynamics across a systematic grid of the movement kernel and epidemic growth rate space to confirm that the simulator achieved a wide range of epidemic structures (Figure S3). As anticipated, epidemics spread to many cells, quickly in unmanaged scenarios with high $\beta$-values. This growth occurred as a clear propagating process away from the location of the index cases when host movement propensities were low (which is to say, $c$-values in the dispersal kernel function were high), but spread rapidly throughout the entire simulation space when host movement propensities were high (which is to say, $c$-values were low).

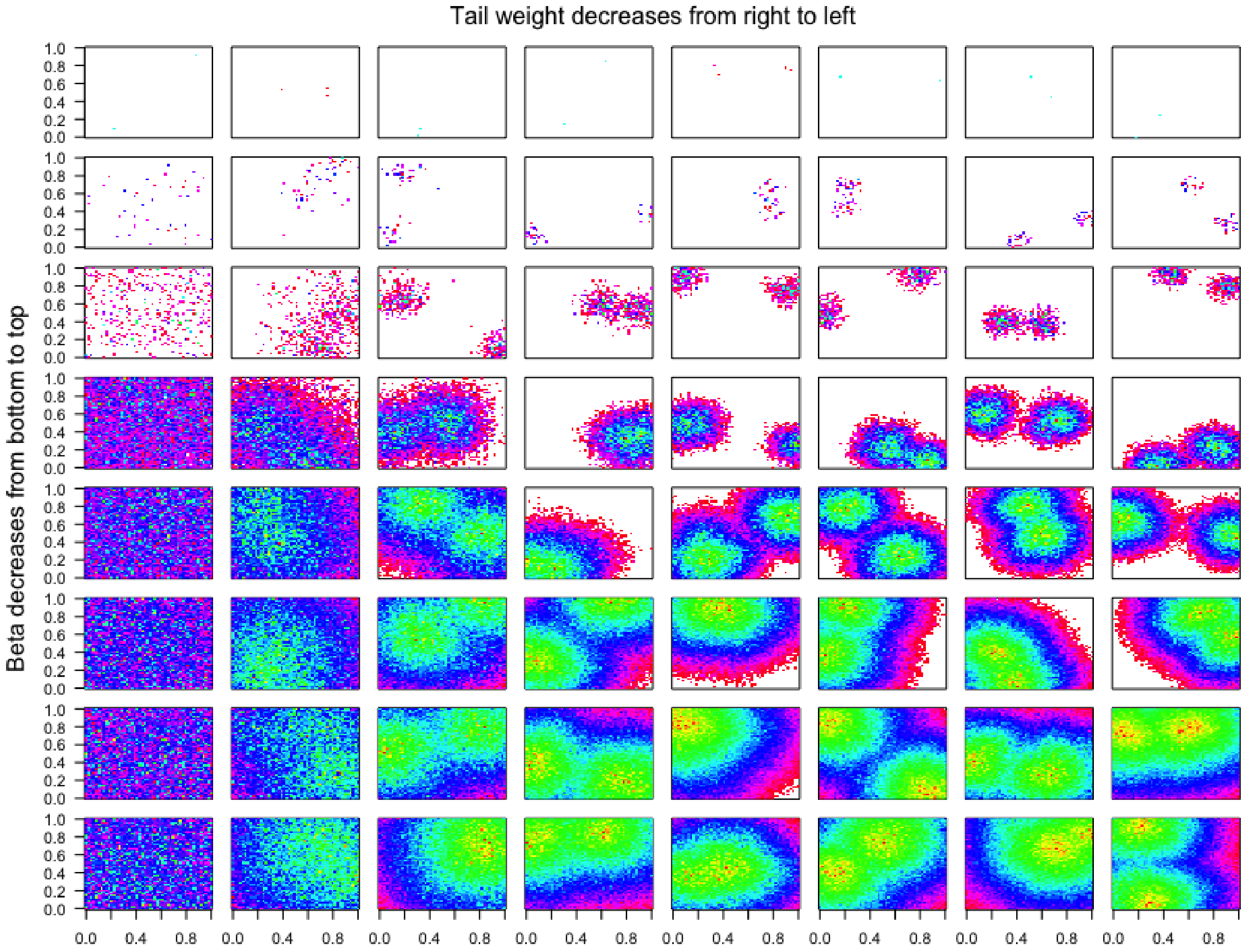

Figure S3: Spatial maps of simulated pandemics in the reservoir host, in the absence of management. Dimensions within individual plots are two-dimensional pictures of space (i.e., "latitude" and "longitude"). Cell colors reflect the first timestep that cell was infected, with the center red point being the initiating cells, ranging through yellow (earliest) to fuschia (last infected). .

We quantified the spatial configuration of each epidemic within the reservoir host more explicitly to be sure spatial propagation was operating as intended. At the end of the simulation, we categorized all cells as ever experiencing infection during the simulation (cells were assigned a 1 if they became infected at any time during the simulation, and 0 otherwise). We then removed all uninfected cells, and constructed a spatial network of infected cells, in which cells were connected to their direct spatial neighbors. This gave us a transmission network (albeit an undirected one). We calculated the number of components (isolated groups of contiguous, infected cells) and the maximum component size in the transmission network to determine how many isolated patches became infected over the course of the epidemic. Number of components gave us a coarse metric of the epidemic's fragmentation over the landscape. This same protocol was also adopted for recipient host cells. We then examined propagation dynamics in the reservoir to be sure that the epidemic was more likely to create a giant connected component when $c$ was high (i.e., host movement propensities were low), but disaggregated into multiple small epicenters of infection when $c$ was low (i.e., host movement propensities were high; Figure S4).

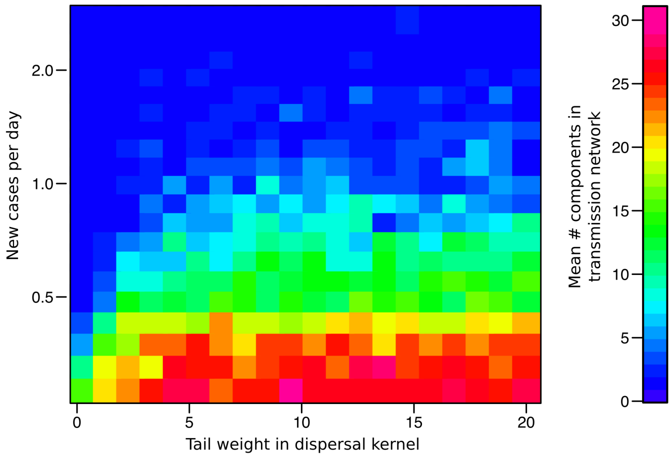

Figure S4: Phase transitions in wave dynamics. Transmission modulates between occurrence within a giant connected component (blue) and occurrence across dispersed, spatially disjoint epidemics (red) over our two-dimensional parameter space in the reservoir host. Consistent with previous work [21], the patterns indicate that the threshold value for widespread spatial transmission regularly exceeds the conventionally accepted $R_0 = 1$ threshold. This is because contact processes dominated by local contacts quickly become saturated, so that the assumption a "completely susceptible population" is rapidly invalid [22].

We visually assessed the performance of each management action at a few cross-sections of the disease parameter space. An example containing the visualization for high-$R_0$ ($\beta = 4.825$) and moderate $c$ are shown in the Main Text (Figure 2), but we show dynamics under a more complete cross section of epidemic growth rates and movement propensities in Figure S3.

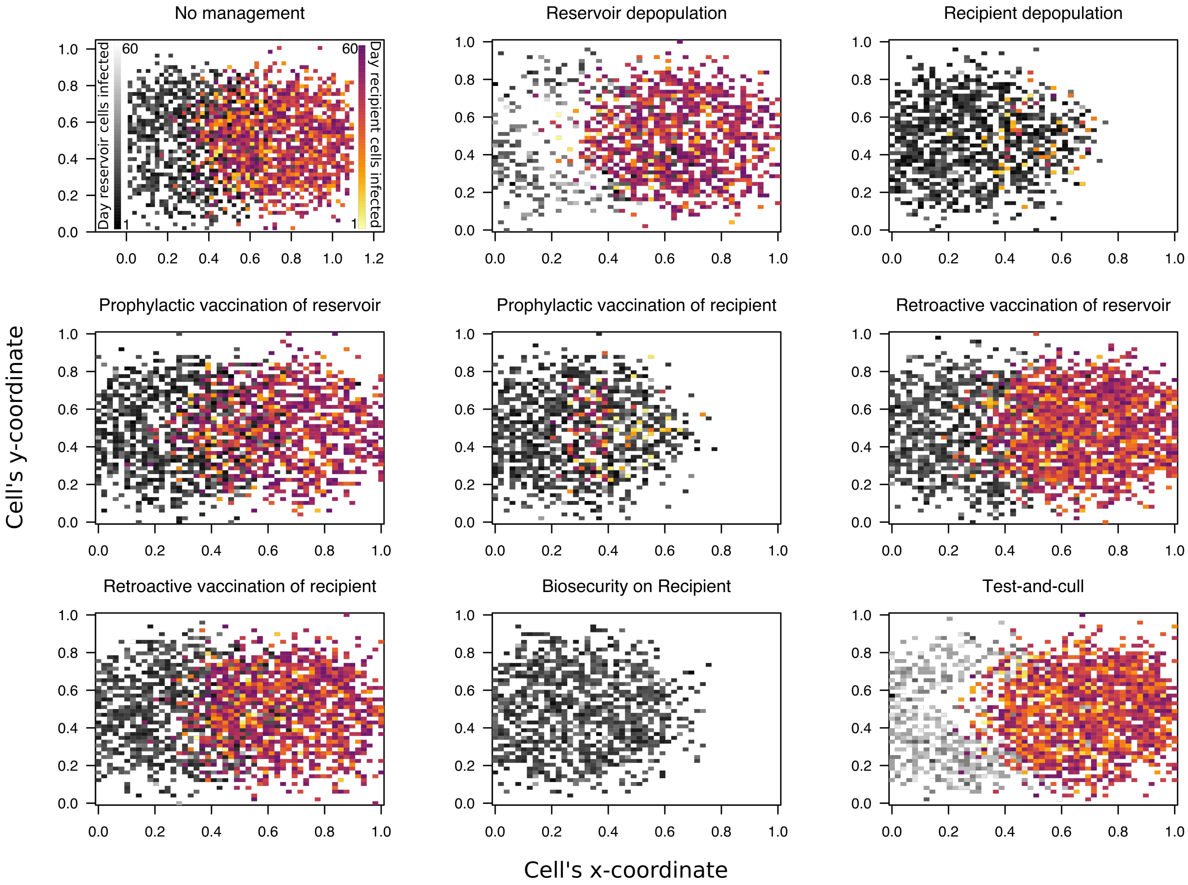

Figure S5: Different management actions simulated on a common disease propagation space.

### 5 Additional results

#### 5.1 Aggregate performance of the management actions

Table 3 shows aggregate performance of each management actions across the entire disease propagation space.

| Objective | Action | % "best" |
|---|---|---|
| Minimise recipient patches | Biosecurity | 0.19 |
| | Depopulation of the Recipient | 0.05 |
| | Depopulation of the Reservoir | 0.07 |
| | Prophylactic Vaccination of the Recipient | 0.06 |
| | Prophylactic Vaccination of the Reservoir | 0.13 |
| | Retroactive vaccination of the Reservoir | 0.19 |
| | Retroactive vaccination of the Recipient | 0.19 |
| | Containment | 0.06 |
| | Selective removal of the reservoir | 0.05 |
| Minimise recipient prevalence | Biosecurity | 0.18 |
| | Depopulation of the Recipient | 0.07 |
| | Depopulation of the Reservoir | 0.07 |
| | Prophylactic Vaccination of the Recipient | 0.07 |
| | Prophylactic Vaccination of the Reservoir | 0.14 |
| | Retroactive vaccination of the Reservoir | 0.18 |
| | Retroactive vaccination of the Recipient | 0.18 |
| | Containment | 0.06 |
| | Selective removal of the reservoir | 0.05 |
| Minimise reservoir patches | Biosecurity | 0.01 |
| | Depopulation of the Recipientp | 0.01 |
| | Depopulation of the Reservoir | 0.05 |
| | Prophylactic Vaccination of the Recipient | 0.02 |
| | Prophylactic Vaccination of the Reservoir | 0.55 |
| | Retroactive vaccination of the Reservoir | 0.26 |
| | Retroactive vaccination of the Recipient | 0.01 |
| | Containment | 0.01 |
| | Selective removal of the reservoir | 0.09 |
| Minimise reservoir prevalence | Biosecurity | 0.15 |
| | Depopulation of the Recipient | 0.15 |
| | Depopulation of the Reservoir | 0.06 |
| | Prophylactic Vaccination of the Recipient | 0.17 |
| | Prophylactic Vaccination of the Reservoir | 0.01 |
| | Retroactive vaccination of the Reservoir | 0.02 |
| | Retroactive vaccination of the Recipient | 0.13 |
| | Containment | 0.15 |
| | Selective removal of the reservoir | 0.15 |

Table S3: Proportion of time each management action performed "best" under each objective, across the entire range of epidemic growth rates and host movement propensities explored here.

#### 5.2 Logistic regression model fits

Table S4: Biosecurity performances

| | *Dependent variable:* | | | |
|---|---|---|---|---|
| | I(Biosecurity was most effective management action) for each of the following: | | | |
| | (Min. recipient patches) | (Min. recipient prevalence) | (Min. reservoir patches) | (Min. reservoir prevalence) |
| ln($\beta$) | | 0.079 | -0.116 | -0.027 |
| | (0.074) | (0.074) | (0.624) | (0.193) |
| ln($c$) | | | 0.065 | 0.234 |
| | (0.068) | (0.067) | (0.483) | (0.180) |
| ln($\beta$):ln($c$) | | | 0.315 | 0.095 |
| | (0.064) | (0.063) | (0.528) | (0.157) |
| Constant | | | | |
| | (0.079) | (0.078) | (0.564) | (0.218) |
| Observations | 1,365 | 1,418 | 126 | 220 |
| Log Likelihood | -546.496 | -558.162 | -17.450 | -80.997 |
| Akaike Inf. Crit. | 1,100.993 | 1,124.325 | 42.899 | 169.995 |

*Note:* Columns contain coefficient estimates (standard errors) for each coefficient in the model corresponding to the column's label.
$^{*}$p<0.1; $^{**}$p<0.05; $^{***}$p<0.01

Table S5: Retroactive vaccination performances

| | *Dependent variable:* | | | |
|---|---|---|---|---|
| | I(Retroactive vaccination was most effective management action) for each of the following: | | | |
| | (Min. recipient patches) | (Min. recipient prevalence) | (Min. reservoir patches) | (Min. reservoir prevalence) |
| ln($\beta$) | | 0.076 | -0.645 | |
| | (0.074) | (0.074) | (0.543) | (1.537) |
| ln($c$) | | | 0.126 | -0.828 |
| | (0.068) | (0.067) | (0.359) | (1.616) |
| ln($\beta$):ln($c$) | | | 0.073 | -0.791 |
| | (0.064) | (0.063) | (0.462) | (1.177) |
| Constant | | | | |
| | (0.079) | (0.078) | (0.426) | (2.108) |
| Observations | 1,365 | 1,418 | 126 | 220 |
| Log Likelihood | -548.188 | -559.934 | -23.271 | -31.717 |
| Akaike Inf. Crit. | 1,104.376 | 1,127.869 | 54.543 | 71.434 |

*Note:* Columns contain coefficient estimates (standard errors) for each coefficient in the model corresponding to the column's label.
$^{*}$p<0.1; $^{**}$p<0.05; $^{***}$p<0.01

Table S6: Prophylactic vaccination (reservoir) performances

| | *Dependent variable:* | | | |
|---|---|---|---|---|
| | I(Prophylactic vaccination was most effective management action) for each of the following: | | | |
| | (Min. recipient patches) | (Min. recipient prevalence) | (Min. reservoir patches) | (Min. reservoir prevalence) |
| ln($\beta$) | | | 0.121 | |
| | (0.066) | (0.063) | (0.208) | (0.141) |
| ln($c$) | 0.071 | 0.048 | 0.026 | 0.098 |
| | (0.059) | (0.056) | (0.169) | (0.131) |
| ln($\beta$):ln($c$) | 0.066 | 0.044 | -0.003 | -0.008 |
| | (0.055) | (0.053) | (0.180) | (0.116) |
| Constant | | | -0.017 | |
| | (0.071) | (0.066) | (0.197) | (0.159) |
| Observations | 1,365 | 1,418 | 126 | 220 |
| Log Likelihood | -740.106 | -771.601 | -87.139 | -120.961 |
| Akaike Inf. Crit. | 1,488.211 | 1,551.201 | 182.277 | 249.923 |

*Note:* Columns contain coefficient estimates (standard errors) for each coefficient in the model corresponding to the column's label.
$^{*}$p<0.1; $^{**}$p<0.05; $^{***}$p<0.01

Table S7: Prophylactic vaccination (recipient) performances

| | *Dependent variable:* | | | |
|---|---|---|---|---|
| | I(Prophylactic vaccination was most effective management action) for each of the following: | | | |
| | (Min. recipient patches) | (Min. recipient prevalence) | (Min. reservoir patches) | (Min. reservoir prevalence) |
| ln($\beta$) | | | -0.045 | |
| | (0.103) | (0.089) | (0.498) | (0.163) |
| ln($c$) | | | -0.235 | 0.007 |
| | (0.091) | (0.079) | (0.408) | (0.152) |
| ln($\beta$):ln($c$) | | 0.118 | 0.239 | -0.047 |
| | (0.083) | (0.074) | (0.434) | (0.134) |
| Constant | | | | |
| | (0.113) | (0.095) | (0.474) | (0.184) |
| Observations | 1,365 | 1,418 | 126 | 220 |
| Log Likelihood | -462.839 | -493.635 | -23.883 | -96.354 |
| Akaike Inf. Crit. | 933.678 | 995.271 | 55.767 | 200.708 |

*Note:* Columns contain coefficient estimates (standard errors) for each coefficient in the model corresponding to the column's label.
$^{*}$p<0.1; $^{**}$p<0.05; $^{***}$p<0.01

Table S8: Reservoir depopulation performances

| | *Dependent variable:* | | | |
|---|---|---|---|---|
| | I(Depopulation of the reservoir was most effective management action) for each of the following: | | | |
| | (Min. recipient patches) | (Min. recipient prevalence) | (Min. reservoir patches) | (Min. reservoir prevalence) |
| $\ln(\beta)$ | | | | |
| | (0.101) | (0.096) | (0.362) | (0.249) |
| $\ln(c)$ | 0.117 | 0.081 | -0.222 | -0.051 |
| | (0.092) | (0.086) | (0.345) | (0.239) |
| $\ln(\beta)$:$\ln(c)$ | 0.057 | 0.047 | -0.228 | 0.013 |
| | (0.085) | (0.080) | (0.312) | (0.205) |
| Constant | | | | |
| | (0.110) | (0.103) | (0.409) | (0.287) |
| Observations | 1,365 | 1,418 | 126 | 220 |
| Log Likelihood | -364.294 | -388.340 | -46.255 | -66.922 |
| Akaike Inf. Crit. | 736.589 | 784.679 | 100.511 | 141.845 |

*Note:* Columns contain coefficient estimates (standard errors) for each coefficient in the model corresponding to the column's label.
$^{*}$p<0.1; $^{**}$p<0.05; $^{***}$p<0.01

Table S9: Recipient depopulation performances

| | *Dependent variable:* | | | |
|---|---|---|---|---|
| | I(Depopulation of the recipient was most effective management action) for each of the following: | | | |
| | (Min. recipient patches) | (Min. recipient prevalence) | (Min. reservoir patches) | (Min. reservoir prevalence) |
| $\ln(\beta)$ | | | | |
| | (0.101) | (0.086) | (0.495) | (0.165) |
| $\ln(c)$ | | 0.094 | -0.045 | 0.061 |
| | (0.089) | (0.076) | (0.310) | (0.154) |
| $\ln(\beta)$:$\ln(c)$ | | 0.085 | 0.341 | 0.066 |
| | (0.082) | (0.072) | (0.416) | (0.136) |
| Constant | | | | |
| | (0.110) | (0.091) | (0.370) | (0.186) |
| Observations | 1,365 | 1,418 | 126 | 220 |
| Log Likelihood | -469.663 | -503.414 | -29.918 | -95.573 |
| Akaike Inf. Crit. | 947.325 | 1,014.828 | 67.835 | 199.146 |

*Note:* Columns contain coefficient estimates (standard errors) for each coefficient in the model corresponding to the column's label.
$^{*}$p<0.1; $^{**}$p<0.05; $^{***}$p<0.01

Table S10: Reservoir selective removal performances

| | *Dependent variable:* | | | |
|---|---|---|---|---|
| | I(Selective removal was most effective management action) for each of the following: | | | |
| | (Min. recipient patches) | (Min. recipient prevalence) | (Min. reservoir patches) | (Min. reservoir prevalence) |
| ln($\beta$) | | | -23.835 | |
| | (0.125) | (0.109) | (3,550.751) | (0.226) |
| ln($c$) | | 0.101 | -10.241 | 0.070 |
| | (0.111) | (0.099) | (1,578.818) | (0.216) |
| ln($\beta$):ln($c$) | 0.085 | 0.025 | -14.629 | 0.010 |
| | (0.102) | (0.092) | (2,206.206) | (0.186) |
| Constant | | | -19.767 | |
| | (0.135) | (0.118) | (2,541.010) | (0.260) |
| Observations | 1,365 | 1,418 | 126 | 220 |
| Log Likelihood | -283.922 | -316.200 | -11.636 | -76.913 |
| Akaike Inf. Crit. | 575.845 | 640.401 | 31.273 | 161.827 |

*Note:* Columns contain coefficient estimates (standard errors) for each coefficient in the model corresponding to the column's label.
*p<0.1; **p<0.05; ***p<0.01

Table S11: No management performances

| | *Dependent variable:* | | | |
|---|---|---|---|---|
| | I(No action was most effective management action) for each of the following: | | | |
| | (Min. recipient patches) | (Min. recipient prevalence) | (Min. reservoir patches) | (Min. reservoir prevalence) |
| ln($\beta$) | | | -0.148 | 0.118 |
| | (0.104) | (0.093) | (0.518) | (0.197) |
| ln($c$) | | | -0.034 | |
| | (0.092) | (0.082) | (0.401) | (0.182) |
| ln($\beta$):ln($c$) | | | 0.422 | -0.151 |
| | (0.084) | (0.076) | (0.438) | (0.159) |
| Constant | | | | |
| | (0.113) | (0.100) | (0.469) | (0.222) |
| Observations | 1,365 | 1,418 | 126 | 220 |
| Log Likelihood | -462.510 | -487.198 | -23.562 | -80.613 |
| Akaike Inf. Crit. | 933.020 | 982.395 | 55.124 | 169.227 |

*Note:* Columns contain coefficient estimates (standard errors) for each coefficient in the model corresponding to the column's label.
*p<0.1; **p<0.05; ***p<0.01

#### 5.3 Fits under other objectives

Figures S6 through S8 show results parallel to Figure 4 in the main text, but for the other three output metrics: recipient prevalence; reservoir patches infected, and reservoir prevalence.

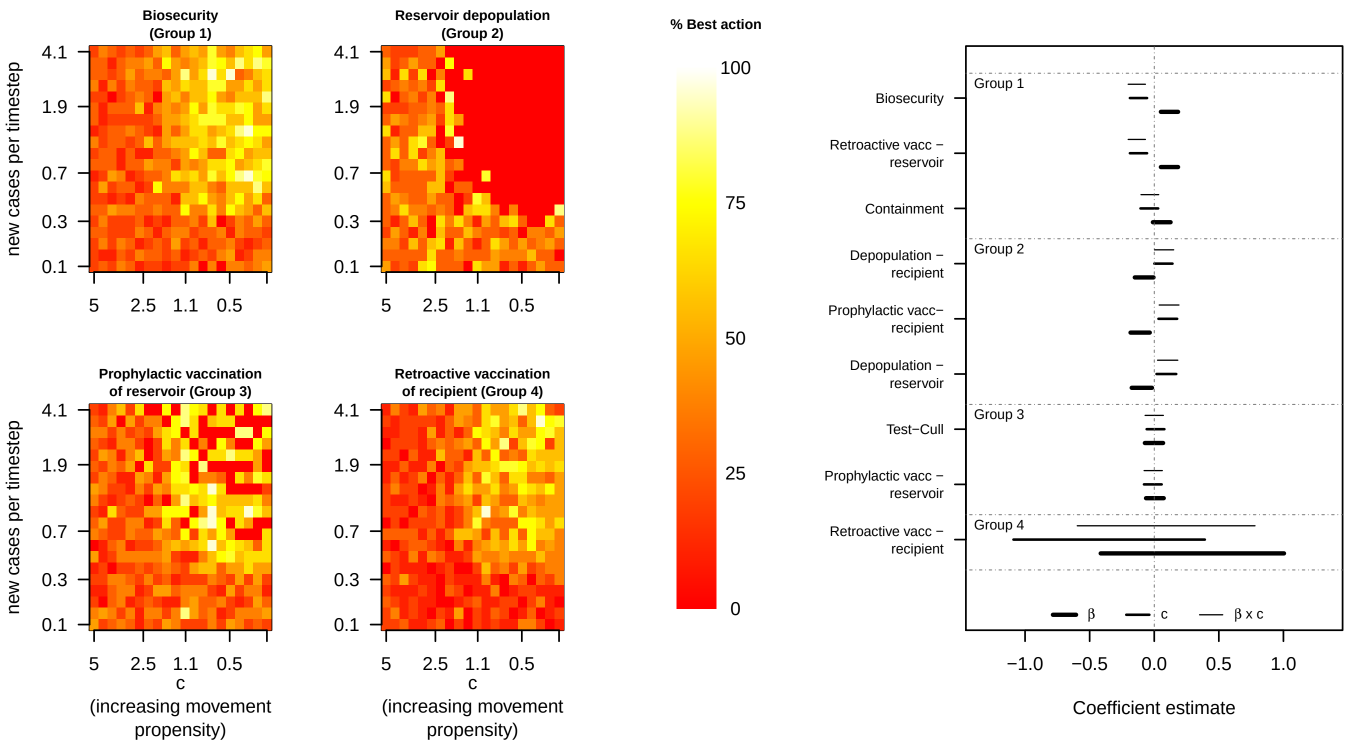

Figure S6: Relative management performance and model coefficient estimates when the response metric was the number of recipient patches infected.

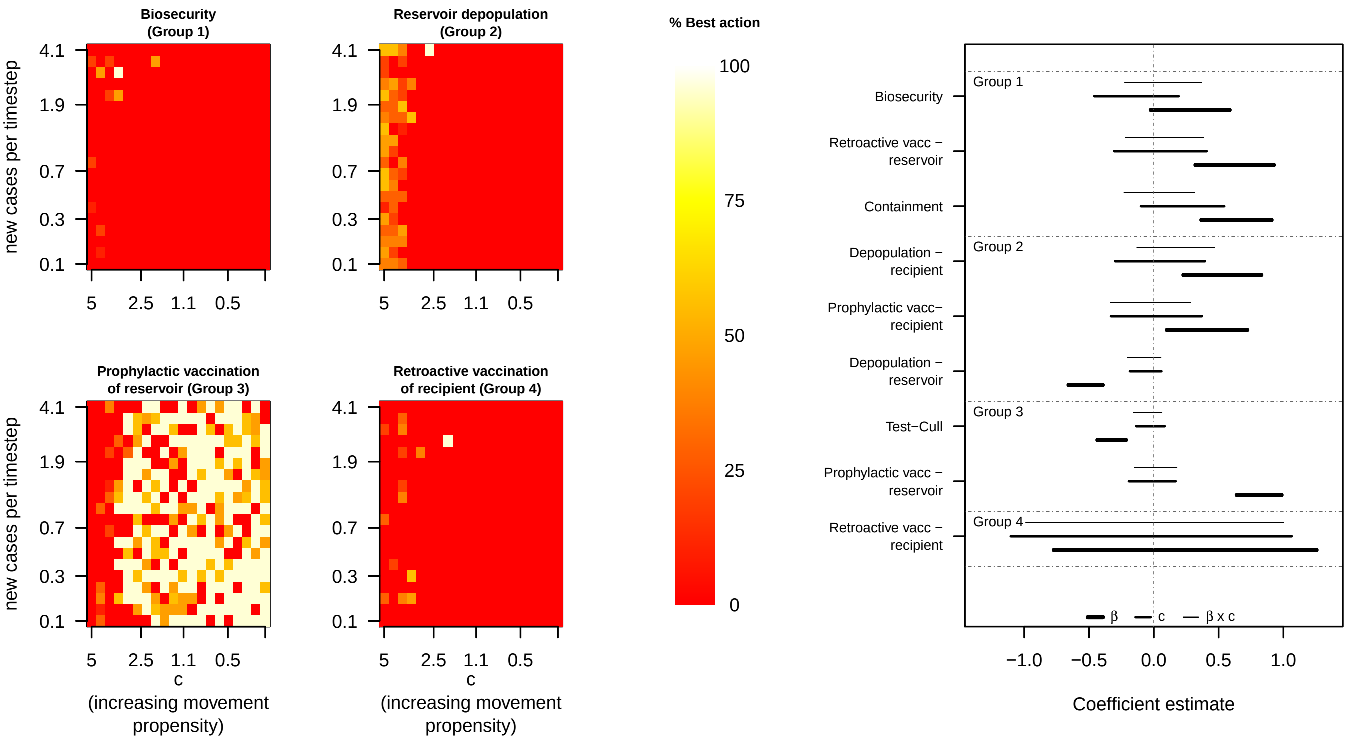

Figure S7: Management competition and model coefficient estimates when the response metric was total reservoir patches infected.

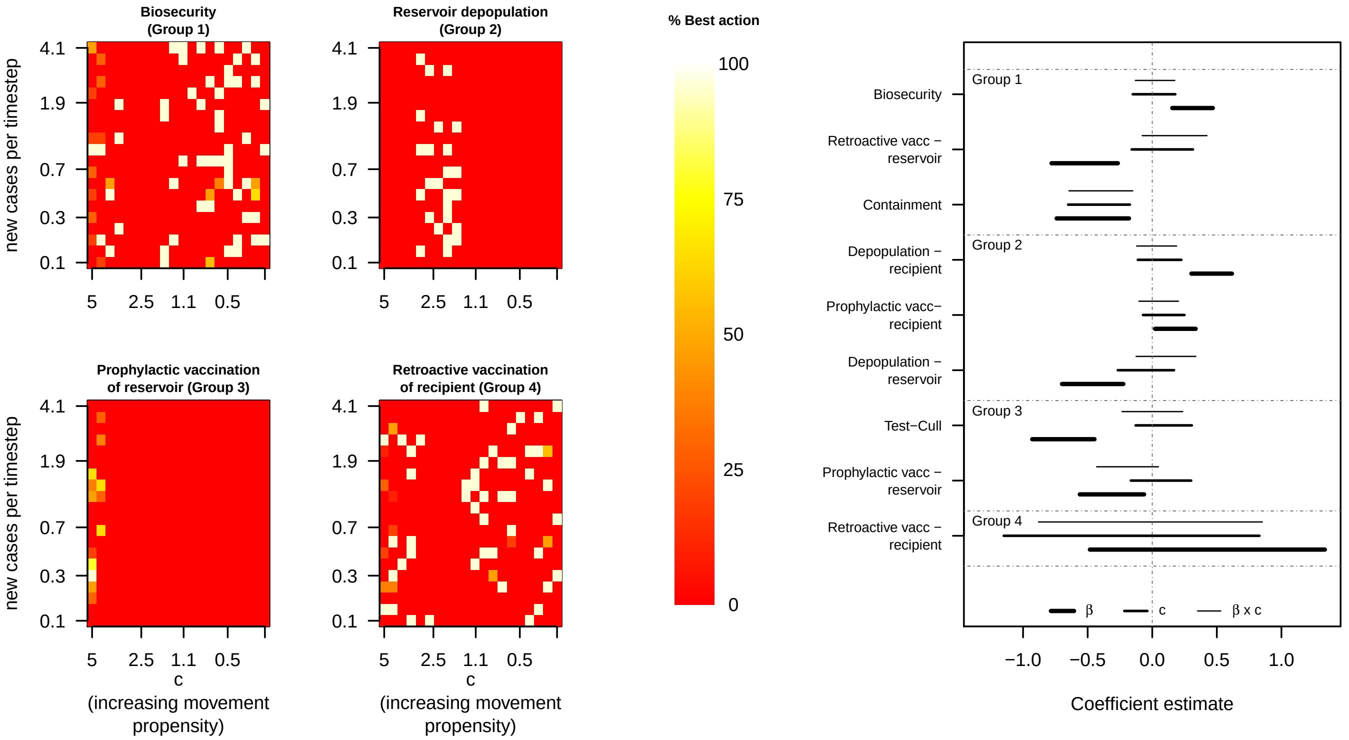

Figure S8: Management competition and model coefficient estimates when the response metric was aggregate prevalence in the reservoir host.

#### 5.4 Classification tree approach

In addition to the logistic regression models, we also used a classification tree to to assess the role epidemic growth rate and host movement propensities played in shaping optimal management within a context where we also considered variation in spatial divide between host species, prevalence that triggered management to start, and reservoir host population densities. We fit four regression trees [23] with response values corresponding to each of our objectives (i.e., total number of recipient patches infected; maximum recipient prevalence, total number of reservoir patches infected to identify the most effective management action according to information on all six covariates. Briefly, regression trees operate by assuming a constant response model within a specified partition of the covariate space. The objective of tree-based methods is to define a path of binary splits that optimises that minimises variation in the response variable within partitions, while maximising variance among partitions. In our case, this equated to identifying covariate values at which the objective function's measured value changed substantially. The size of the trees — which is to say, the number of partitions — governs the model's complexity. We followed standard protocols of growing a very large tree, and then pruning it back to include only splits up to and including the split that minimised cross-validation error. Tree partitioning was implemented using the `rpart` package in `R` [24].

Recursive partitioning methods identify a progressive set of covariate values that best split a set of varying outcomes into groups. Once a partition is identified, subsequent partitions operate exclusively within existing groups (so that the second partitioning of one group might rely on a different covariate than the second partitioning of a different group). Once the outcomes are completely partitioned, the resulting binary tree is pruned back via cross-validation to appropriately avoid overfitting.

We used the Gini impurity criterion for the classifier, with data weights proportional to the observed frequencies of each treatment combination (this was very nearly balanced in the dataset, since we controlled the simulation's parameter space). Any risk within one standard error of the achieved minimum is marked as being equivalent to the minimum (i.e. considered to be part of the flat plateau). Then the simplest model, among all those "tied" on the plateau, is chosen.

Classifier performance was evaluated through cross-validation and trees were pruned to the complexity level associate with the minimum cross-validation error. We fit separate trees for each of four objective functions (minimizing spatial extent or prevalence in the recipient or reservoir host). Pruned trees, along with variable importance estimates in each case, are shown in Figure S7.

Variable importance from the four regression trees consistently indicated that epidemic growth rate and host movement propensities were the most important factors in determining epidemic size and spatial extent, especially when objective functions focused on the recipient host (Figure S8). Spatial separation of reservoir and recipient host activity centers and management actions were also important determinants of epidemic size and extent.

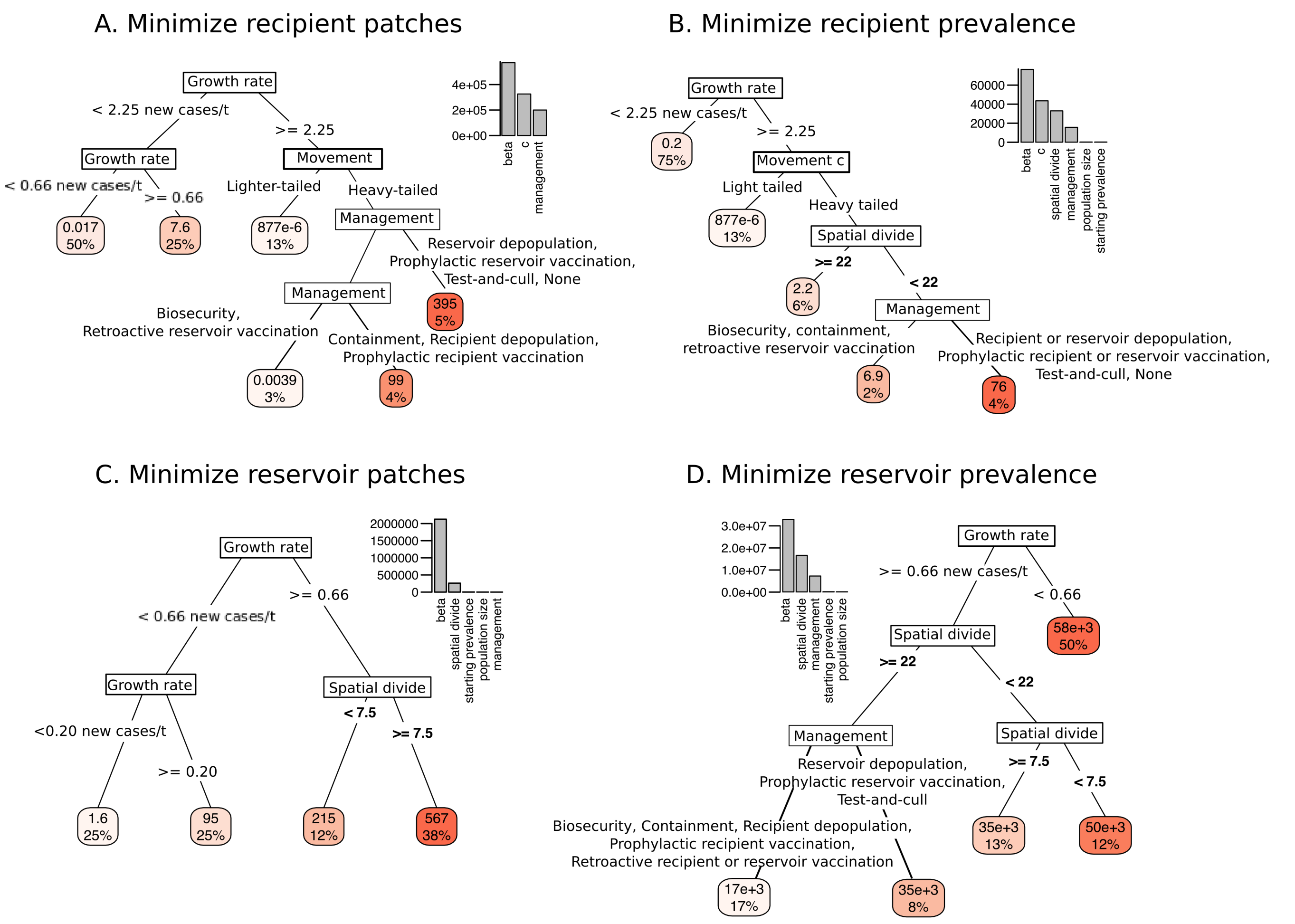

Figure S9: Regression trees showing process and management parameters associated with varying values of each of four objective functions. Leaf colors represent epidemic size (in terms of patches or prevalence), with redder leaves being larger epidemics in the specified metric. Leaf percentages reflect the total proportion of simulations landing in each leaf.

### 6 Limitations associated with this framework

#### 6.1 SIR assumptions and limitations

First, commensurate with our SIR modeling structure, we assumed that any pathogen infection provided hosts with complete immunity to that pathogen in the future. However, we know this assumption is violated in several key wildlife-livestock spillover diseases (including avian influenza, leptospirosis, and bighorn sheep pneumonia, to name a few). Accounting for partial or limited cross-strain immunity would likely have slowed reservoir fade-out in the fastest-growing cases (but probably would not have lower prevalence), since epidemics would have had a larger pool of susceptible hosts available. Thus our model probably underestimates reservoir prevalence and subsequent spillover burden for diseases with partial immunity.

Second, we neglected disease-induced mortalities throughout this exploration. This might be a defensible assumption for diseases that we manage foremostly due to their downstream risks to human health (i.e., brucellosis; North

American rabies), and possibly also for diseases that pose limited consequences on reservoir host health (i.e., rabies in bats; M. ovi in domestic sheep). Limited disease induced mortality is inconsistent with many diseases of management concern at the wildlife-livestock interface. Disease-induced mortality would likely lower spillover risk in many systems for two reasons. First, mortalities curtail the duration of the infectious period, and may limit the movement potential of infected animals. Second, ill animals may be less-likely to move than their infected counterparts.

Densities would also be altered by disease-induced mortalities, and even beyond disease-related changes, many — if not most— wildlife and livestock systems in temperate latitudes exhibit seasonally pulsed densities. Varying densities would introduce additional variation into transmission rates for pathogens with density-dependent transmission routes. Decreases, and even simply oscillations in densities are thought to to drive pathogens toward local extinction however (Peel et al. 2014), so our choice to create densities as constant likely biases our model toward over-estimating spillover frequencies.

We took process parameters (per-susceptible transmission rate, recovery rate, movement rate) to be constant. but these could also feasibly change over the course of a spillover event. In the most basic case, some hosts have fundamentally different transmission parameters than others due to switches in mode of transmission that co-occur with host shifts (for instance, avian influenza's switch from primarily gastrointestinal to primarily respiratory when it switches from wild to domestic fowl). Human-mediated movement dynamics (as is especially common in livestock hosts) almost certainly change once a spillover event is detected and reported, with strong consequences on post-spillover epidemic growth rates.

#### 6.2 Timescale and epidemic duration

We chose 60 timesteps as the duration for all simulations. This, and any other, timescale choice is somewhat arbitrary, since both epidemic dynamics and spatial movements accumulate continuously in time (though we update movements in batches; see Supplementary Materials: tau-leap). However, the 60-timestep scale aligned with our transmission, recovery, and movement rates to provide a wide range of epidemic dynamics. Additionally, it seemed reasonable that management agencies might be able to categorize pathogens as expanding on a weekly (i.e., 1 timestep), seasonal (i.e., 12 timestep), or annual (i.e., 52 timestep) scale, and to implement some management responses at a weekly scale, but probably not much faster.

#### 6.3 Direction and independence of movements

Our simulation landscape had no structure beyond cell-to-cell distance, so distance was the only determinant of where individuals moved. Also, since we held all within-cell populations constant, there was no "crowding" effect. These assumptions are clearly violated at some level for most real-world animal systems. We also force animals to move as independent units, overlooking larger-scale migrations or group-level moves. This likely means that the number of independent movers is biased high in our simulations, but the capacity of those movers to spark an epidemic could be biased low (since if individuals actually move in groups of 5, for example, any one of the five movers could be infected and spark an epidemic). However, without information specific to the behavioral ecology and spatial context of a given host system, we felt that adding additional detail here likely caused more problems than it alleviated.

#### 6.4 Common movement kernels for reservoir and recipient hosts

In this simulation, we assumed that both reservoir and recipient host species moved according to identical movement kernels. This assumption was made for the purposes of simplicity, and is unlikely to hold in many wildlife-livestock situations. There are, however, a few places where it could be appropriate, and we highlight those instances here.

One context where common kernels could be reasonable is for host species that are closely related or allometrically matched (for instance, a system in which both the reservoir and the recipient host species are ungulates; a system where both hosts are canids, etc.), and both experience largely uninhibited movements (on the livestock side, this could include livestock that are ranged on grazing allotments, or animals like free-ranging domestic cats and dogs living at the urban-wildland interface).
